# COCOA-Tree: Phylogenetic visualization and comparative analysis of coevolving residues

**DOI:** 10.64898/2026.02.05.703816

**Authors:** Margaux Jullien, Marion Chauveau, Sophie-Carole Chobert, Emma Bouvet, William Schmitt, Fabien Pierrel, Sophie Abby, Nelle Varoquaux, Ivan Junier

## Abstract

The evolutionary co-occurrence of amino acid changes between protein residues underlies key structural and functional properties of protein families. Building on these coevolutionary patterns, methods have been developed to identify groups of residues associated with enzyme functionalities, such as Statistical Coupling Analysis (SCA) or Specificity-Determining Position (SDP) methods. These methods and their variations differ in the metrics used to quantify coevolution, residues weighting schemes, and corrections introduced to mitigate noise and phylogenetic biases. Yet, systematic comparisons across methods are rarely performed, and the evolutionary origins of the coevolution patterns highlighted by each approach are seldom addressed, limiting our ability to disentangle functional from phylogenetic contributions.

To address these issues, we introduce COCOA-Tree, a Python library for SCA-like dimensionality-reduction analyses. COCOA-Tree supports custom metrics and enables visualization of coevolutionary patterns on phylogenetic trees. We also provide guidance to map results onto 3D structures in PyMOL. Using COCOA-Tree, we reanalyze published datasets and uncover previously unnoticed evolutionary properties of groups of coevolving residues detected by SCA, known as sectors. In particular, in the well-studied S1A serine protease family, we show that two of the three known sectors exhibit qualitatively distinct levels of sequence conservation depending on the enzymatic functions and on the phylogenetic clades to which the proteins belong. We further show that different coevolution metrics often identify qualitatively distinct groups of coevolving residues, although they yield consistent results for mildly conserved residues. Finally, as an example of the versatility of COCOA-Tree, we provide an example of visualizing pairs of residues detected by Direct Coupling Analysis (DCA) methods on a phylogenetic tree, highlighting a rich diversity of co-evolutionary patterns.

Overall, we expect COCOA-Tree to help identify residues that control protein function and thereby improve our capacity for functional engineering and our understanding of the principles governing protein evolution.

COCOA-Tree website: https://tree-timc.github.io/cocoatree

## Introduction

Coevolutionary analyses of protein residues underlie the fields of protein structure prediction and enzyme-function rationalization [1–3]. The general idea behind these methods is to identify coevolving residues by analyzing a multiple sequence alignment (MSA) of a protein family, which can then be used to infer structural contacts and functional determinants. Technically, the dimensionality reduction of the original MSA to a smaller set of residues and relationships can be achieved in multiple ways, depending on both the problem to be solved and the methodology adopted. In this regard, methods can be distinguished according to whether they aim to detect pairs of co-evolving residues, as in Direct Coupling Analysis (DCA) [4], or larger groups of co-evolving residues. In the latter context, Statistical Coupling Analysis (SCA) seeks to identify so-called sectors associated with protein functions [5] by applying dimensionality reduction to a covariance-like matrix in which residues are weighted according to their amino-acid conservation [6]. In contrast, methods aiming to determine Specificity-Determining Positions (SDPs) often quantify coevolution trends using mutual-information-based metrics [1, 7]. In addition, several corrections such as the Average Product Correction [8] have been developed to mitigate noise arising from limited sequence numbers, or confounding phylogenetic effects [8–13].

The existence of different coevolution metrics, diverse correction schemes, and various residue-weighting procedures complicates the systematic comparison of results and the interpretation of the evolutionary origins of identified coevolving groups [12, 14, 15]. Altogether, this raises fundamental questions: Which methodological choices are most appropriate for identifying coevolving groups? How do the evolutionary properties of these groups relate to those of the rest of the protein sequence? Along these lines, we also note that methods such as SCA do not use phylogenetic relationships, whereas SDP-based methods are intrinsically phylogenetic. Namely, SDPs are defined as “groups of positions that coordinately mutate in the context of subfamily [phylogenetic] divergence” [1]. Finally, these questions, and particularly the role of phylogeny [16], also pervade the problem of detecting specific pairs of co-evolving residues. For instance, multiple tools have been developed to account explicitly for the phylogenetic relationships among sequences [17–20] or to mitigate their effects through phylogeny-aware reshuffling methods [13, 21]. We note, nevertheless, that none of these methods provides a tool, either as a web server or a library, for visualizing the resulting co-evolutionary signals on a phylogenetic tree.

In this context, we present COCOA-Tree, a Python library that integrates SCA-like dimensionality-reduction methods with phylogenetic and metadata visualization to help elucidate the origins of coevolution and its relationship to enzyme function. Specifically, COCOA-Tree allows users to (i) easily generate coevolution matrices under various metrics, corrections, and weighting schemes, (ii) perform dimensionality-reduction analyses and identify associated groups of coevolving residues as in SCA, and (iii) visualize results along phylogenetic trees in the spirit of SDP methods. In addition, we provide guidance for exporting outputs to PyMol for 3D structural mapping. As an illustrative example of the value of COCOA-Tree, we revisit the well-studied S1A serine-protease family and uncover previously unnoticed properties of sector-associated sequence evolution. We also discuss, for this family as well as for two other well-characterized protein families – dihydrofolate reductases (DHFRs) and rhomboid proteases – the impact of coevolution metrics and corrections on the results. Finally, we show an example of visualizing, on a phylogenetic tree, DCA pairs of co-evolving residues identified using adabmDCA, an adaptive Boltzmann-machine learning framework [22], revealing a rich diversity of co-evolutionary patterns.

## Approach

Just as in SCA analyses, one of the main outcomes of COCOA-Tree for a given coevolution metric, correction, and residue weighting scheme is a set of groups of residues obtained by applying a dimensionality reduction method to the corresponding coevolution matrix, more specifically independent component analysis (ICA). In SCA, the resulting “independent” components (ICs)^1^ can be further fused on the basis of the remaining level of correlation between components [6], leading to the biologically motivated notion of *sectors*. Specifically, Ranganathan and co-workers have argued that sectors provide a functional decomposition of proteins associated with independent modes of selection [5, 6, 24]. Here and in the COCOA-Tree toolbox, we focus on ICA results and do not address the fusion of components, as this lies beyond the scope of the tool and requires careful biological interpretation.

Following the SCA methodology and adapting the PySCA code (Methods), we thus define for each IC a group of the most extremal residues of each IC, which we refer to as a XCoR (group of e<u>X</u>tremal <u>Co</u>-evolving <u>R</u>esidues). A XCoR is therefore a group of residues that contribute the most to a given IC, and a XCoR sequence is the sequence obtained by mapping each full-length protein sequence onto only those selected residues. Starting from a MSA, a COCOA-Tree analysis then typically proceeds through the following steps, with steps 1–3 directly adapted from SCA [6]:

1. compute a selected coevolution metric, along with its possible correction and residue weighting,
2. perform a decomposition using ICA,
3. extract the corresponding XCoRs,
4. visualize the XCoRs in the context of phylogenetic trees,
5. and export results for visualization in PyMOL.

Note that the PyMOL export of results enables exploration of the 3D conformations of the proteins and their XCoRs. Coevolving residues, more particularly sectors [5, 6, 24], have indeed been shown to correspond to spatially connected networks of amino acids.

The COCOA-Tree library is divided into four main parts (Fig. 1): i) *data* input and handling, ii) *statistics and metrics*, which allows to compute residue and sequence statistics as well as pairwise statistics underlying coevolution matrices, iii) *decomposition* allowing to perform dimensionality reduction and to extract ICs and XCoRs, and iv) *visualization* of XCoRs in the context of phylogenetics and structural biology.

**Figure 1.**
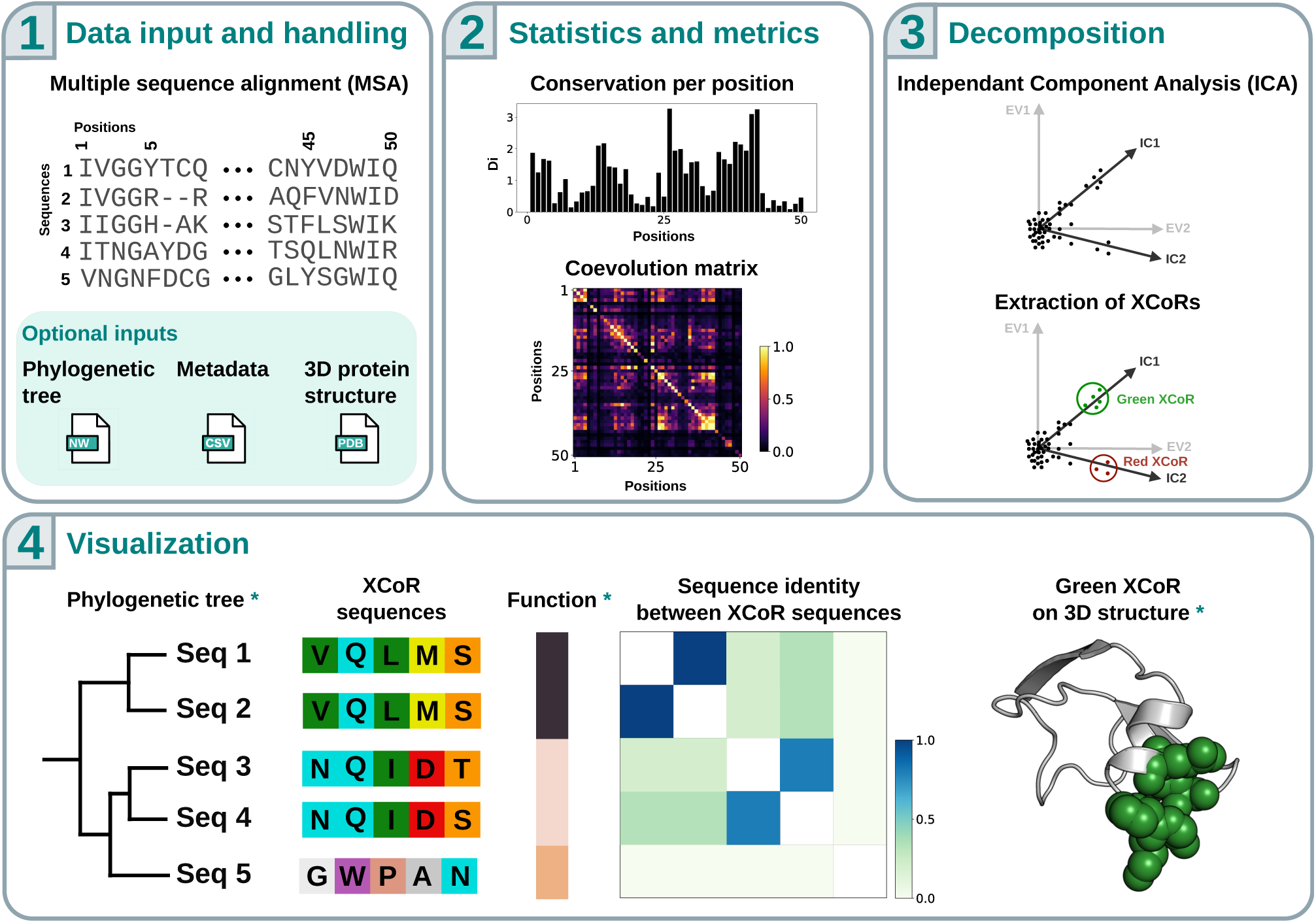
COCOA-Tree’s library general scheme and organization. COCOA-Tree is divided into four main parts. **1.** *Data input and handling*, for loading MSA and corresponding data files. **2.** *Statistics and metrics*, such as the degree of conservation of each residue (or position) and coevolution matrices. **3.** *Decomposition methods*, also referred to as dimensionality-reduction methods, based on an eigenvalue/eigenvector (EV) decomposition followed by ICA, which together yield the identification of XCoRs. In this schematic example, two XCoRs are identified: a green XCoR composed of five residues and a red XCoR composed of three residues. **4.** *Visualization* of the XCoR sequences along phylogenies, as well as PyMOL output files for mapping onto protein 3D structures. Outputs depending on optional inputs (see Panel 1) are marked by green asterisks. In this example, the red XCoR is used as the reference: sequence identity with respect to this XCoR is shown alongside the phylogenetic tree, and its residues are highlighted on the 3D protein structure.

### Data input and handling

COCOA-Tree relies on four types of data (Fig. 1.1): i) a MSA of protein sequences from the family of interest in FASTA format, ii) a phylogenetic tree representing evolutionary relationships between the MSA sequences in Newick format, iii) metadata associated with the MSA sequences in CSV format, and iv) 3D protein structure data for a reference sequence in PDB format. Note that only the MSA data is required. Note also that MSA alignment quality may impact the results and, hence, should be considered a potential variable for the user to explore. Along these lines, several operations can be applied to the MSA in COCOA-Tree, such as filtering overly gapped positions, removing highly gapped sequences, or selecting specific sequences. As examples, COCOA-Tree includes three MSAs for which sectors have previously been identified and analyzed in the context of experimental data or molecular dynamics simulations (see below). The three other input data are optional, as they are used for specific visualizations. Namely, the phylogenetic tree allows visualization of how the identity of the XCoRs varies according to the evolutionary relationships of input protein sequences. The metadata file enables the addition of contextual information, such as taxonomic classification or functional annotations (*e.g.*, enzymatic activity or substrate specificity). The PDB file allows visualization of XCoRs on a representative 3D structure of a selected reference sequence.

COCOA-Tree further includes three datasets from publications that applied the SCA methodology, which are described in more detail in the Methods section: (i) the S1A serine protease dataset of [5], hereafter referred to as the S1A dataset; (ii) the dihydrofolate reductase (DHFR) dataset of [25], which provides recent additional insights relative to the original analysis of [24]; and (iii) the Rhomboid dataset corresponding to a family of intramembrane serine proteases [26]. For each dataset, we provide the MSA as reported in the corresponding article, together with the results of the original SCA method, a phylogenetic tree (Methods), a metadata file mapping each MSA sequence to relevant information for interpretation, and a representative PDB structure.

### Statistics and metrics

Given an MSA, COCOA-Tree computes a range of statistics that can be broadly categorized into first-order and second-order (pairwise) statistics. First-order statistics are calculated for individual residues or sequences, yielding profiles such as residue (or, equivalently, position) conservation. Second-order statistics capture joint variation between pairs of residues and serve as the foundation for identifying groups of residues that share a common evolutionary signal.

#### Position and sequence-wise statistics

First-order statistics associated with residues include amino acid conservation levels – equivalently, the Shannon entropy at each position – and gap frequencies. For sequences, a key first-order statistic is the number of other sequences *m_s_* that are similar to a given sequence *s* – based on a user-defined sequence identity threshold (0.8 as a default value [6]) – which is used to assign each sequence a weight *ω_s_* = 1*/m_s_*. These weights are used to compute all residue frequencies (both first- and second-order) in order to reduce the impact of over-represented, highly similar sequences datasets [6, 27]. In particular, they define the effective number of sequences, *M_eff_* (Methods).

#### Pairwise statistics: coevolution metrics, corrections, and weighting procedures

COCOA-Tree allows one to compute, and hence, compare, various types of coevolution metrics, corrections, and residue weighting (all detailed in Methods). Namely, it includes three types of coevolution metrics: covariance-based (SCA), mutual information (MI), and normalized MI (NMI), as well as two types of corrections (Average Product Correction–APC– and entropy) and one type of residue weighting (SCA weights). In principle, one can combine these elements altogether such as to produce 3 *×* (1 + 2) *×* (1 + 1) = 18 types of coevolution matrices and we provide a unified API to handle them. In addition, COCOA-Tree allows users to implement their own coevolution metric in a straightforward manner.

### Decomposition

In COCOA-Tree, once a coevolution matrix has been estimated, one can apply one of two decomposition methods: (i) a singular value decomposition (SVD), leading to the identification of eigenvectors (Fig. 1.3), as performed in [5]; and (ii) ICA, as proposed by [6]. The number of retained ICs is set to 3 by default as in [6]. However, COCOA-Tree also allows to automatically estimate a relevant number of ICs associated with the spectral properties of the coevolution matrix following the procedure explained in [6]. From the resulting ICs, amino acids can then be assigned to XCoRs, following the statistical criteria proposed in [6] (Methods).

### Visualization: phylogenetic trees and 3D representations

In order to better understand the origins of the XCoRs and to investigate their relationships with both functional and evolutionary aspects of the underlying full sequences, COCOA-Tree includes a phylogeny-based visualization function. To this end, we use the Python library ete3 [28], which enables the plotting of phylogenetic trees together with quantitative metadata (*e.g.*, taxonomic rank or functional annotations) and the amino acid sequences of the XCoRs. COCOA-Tree also allows the display of a heatmap indicating the level of identity between any two XCoR sequences, thereby facilitating comparison of the sequence content of XCoRs in relation to the phylogenetic tree associated with the underlying full sequences. Note here that any tree can be provided as COCOA-Tree input, provided it is in the standard Newick format, so that, in principle, XCoRs can be confronted with any tree-like information associated with the input sequences. In COCOA-Tree’s example gallery, we also provide an example of PyMOL script allowing the user to directly map and highlight the residues of a XCoR on a protein 3D structure.

## Results and discussion

Using COCOA-Tree, we processed and analyzed three previously studied protein families for which publicly available datasets are included in COCOA-Tree’s datasets collection: (1) the S1A protein family, for which the SCA methodology was originally developed and experimentally tested, using the dataset of [5]; (2) the DHFR protein family, in which a sector was shown to control protein allostery [24], using the dataset of [25]; and (3) the rhomboid protein family, in which sector transplantation was shown to convert substrate specificity [26], using the corresponding dataset. For each dataset, we first filtered the MSA to remove redundant sequences (80% identity) and positions with more than 40% gaps, using similarity and gap filtering threshold values from [6]. Table 1 summarizes the statistics for each dataset (number of sequences before and after filtering, number of residues before and after filtering, etc.).

**Table 1.** Summary statistics for the different datasets. For each dataset, we provide the number of sequences, the number of sequences after filtering for gapped sequences, the number of effective sequences, the number of residues before and after filtering for gapped positions, the number of residues affected to a XCoR, and the number of residues affected to XCoRs in the original study.

|  | Serine protease | Rhomboid | DHFR |
| --- | --- | --- | --- |
| Number of sequences | 1470 | 2767 | 4422 |
| Number of sequences after filtering | 1376 | 2767 | 3806 |
| Number of effective sequences | 1053 | 1792 | 3332 |
| Number of residues | 832 | 135 | 802 |
| Number of residues after filtering | 242 | 135 | 156 |
| Number of residues affected to a XCoR | 73 | 42 | 61 |
| Number of residues found in a XCoR in original study | 64 | 40 | 48 |

### Coevolution matrix and XCoR identification

The first step in SCA-like analyses – and in most coevolution analyses – is to compute a coevolution matrix *C* where each entry *C_ij_* represents the degree of coevolution between residues *i* and *j* based on a chosen coevolution metric and residue weighting scheme (Fig. 2A). From this matrix, one can extract a set of XCoRs, which form the basis for the sector decomposition of protein families. The coevolution matrix can then be reduced and reordered accordingly to highlight the tendency of residues belonging to the same XCoR to coevolve more strongly with one another than with residues from other XCoRs (Fig. 2B). Then, an important, yet often overlooked operation, which can be easily performed in COCOA-Tree, consists in removing a global coevolution tendency that affects all residues (and hence all XCoRs) to a similar extent [Fig. 2BC, 6] – technically, it consists in setting to zero the first eigenvalue of the coevolution matrix.

**Figure 2.**
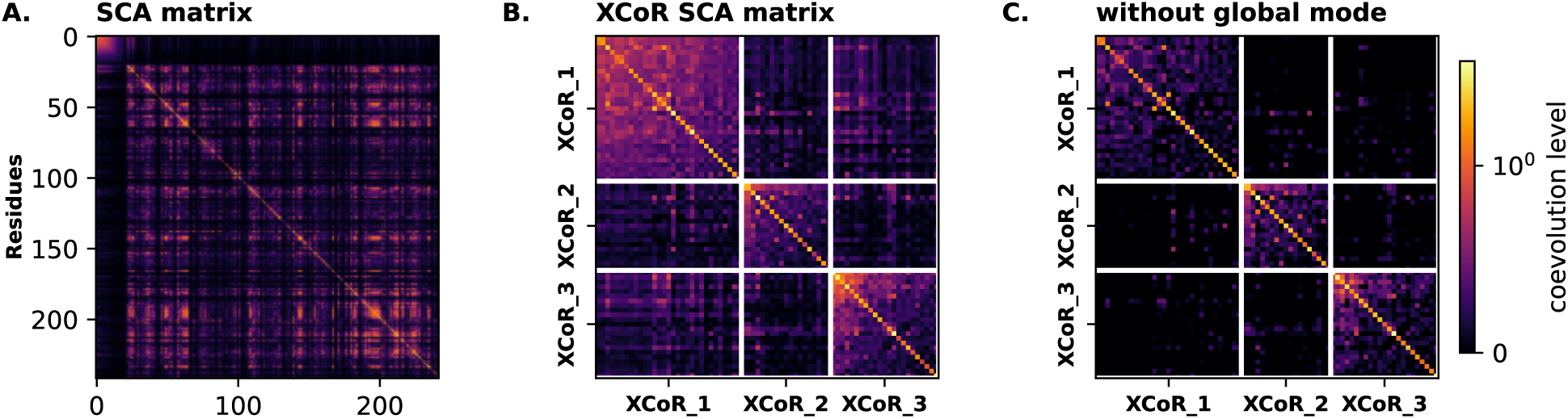
Coevolution matrix sorted according to XCoRs. We display for the S1A dataset the following matrices. **A.** Full SCA coevolution matrix of the 242 MSA positions. **B.** Reduced coevolution matrix of the three sets of XCoRs as defined by an ICA analysis; each white line delimitates an XCoR with residues ordered in decreasing contribution to the corresponding IC. **C.** Coevolution matrix of the three XCoRs without the global mode.

### Reproducing protein sector identification using COCOA-Tree across benchmark datasets

For each dataset, we applied COCOA-Tree using the SCA coevolution metrics – considering the same number of ICs as in the original studies – and assigned residues to the different XCoRs. Because ICA does not have a natural order of components, we identified the best match between COCOA-Tree’s ICs and the original ones based on their XCoR match, and named the components IC_red_, IC_green_, IC_blue_, and IC_purple_ (Fig. 3), with the choice of red, green, and blue matching the sector terminology proposed for the S1A serine protease family [5] – see Table S1 for a mapping between our work’s colors and the other studies.

**Figure 3.**
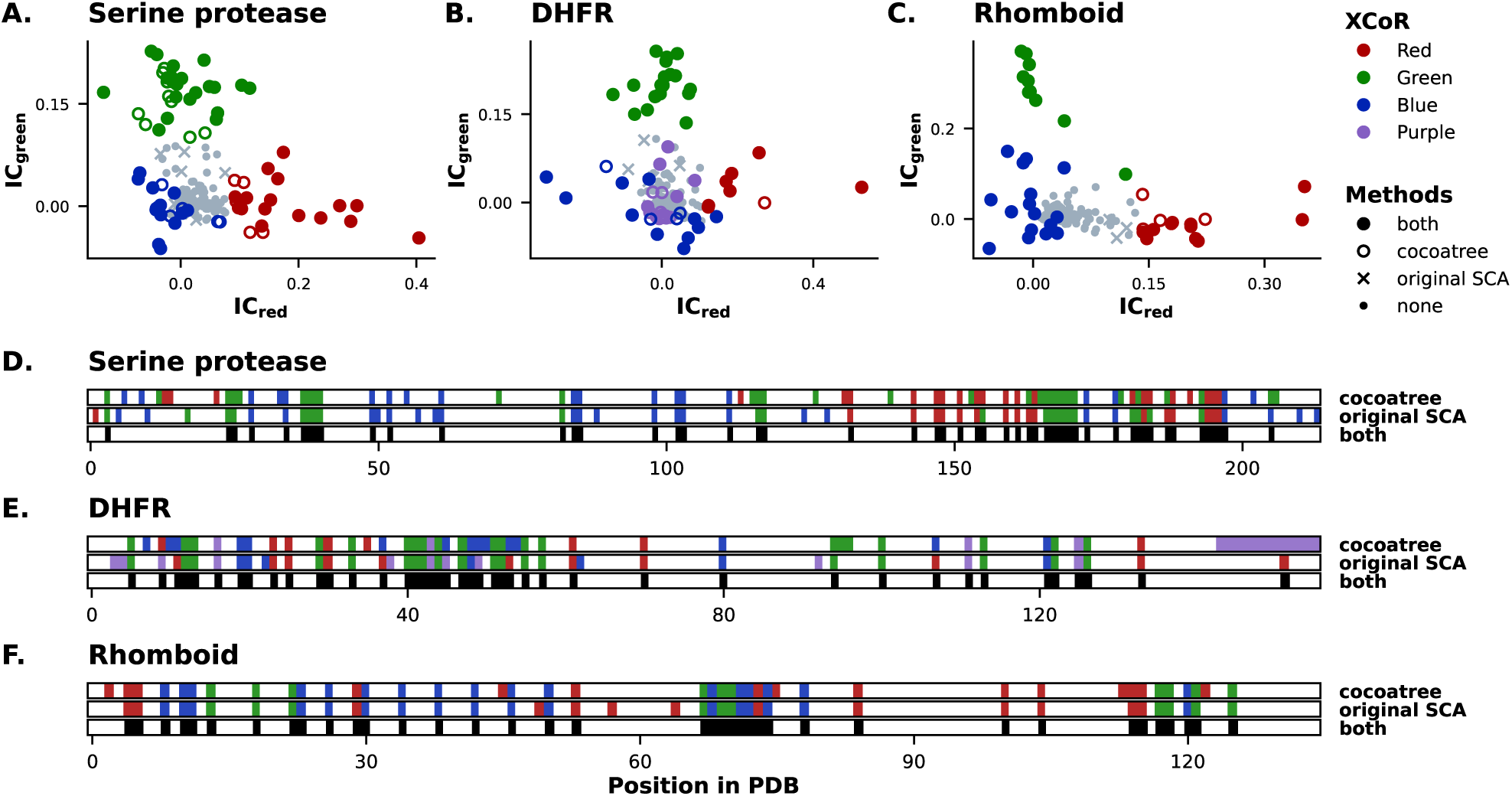
COCOA-Tree XCoRs compared with the “original” ones. ICA performed on **A.** serine protease, **B.** DHFR, and **C.** Rhomboid, showing only the two first components. Colors correspond to XCoRs identified with COCOA-Tree, and markers indicate whether positions are found in both COCOA-Tree and the original analysis, only in COCOA-Tree, only in the original analysis, or in neither tool. **D, E, F** display, for each position in the PDB files of the three datasets, the XCoRs it belongs to according to COCOA-Tree and the original analysis, as well as whether it is shared between them.

Overall, COCOA-Tree finds similar numbers of residues in XCoRs as the original studies (S1A: 64 (our work) versus 73, Rhomboid: 42 vs 40, DHFR: 61 vs 48). The numbers likely differ due to variations in MSA filtering procedures, such as whether positions are filtered before redundant sequences or vice versa. Moreover, examining the residues of each XCoR, we observe substantial overlap for the S1A and rhomboid datasets (Fig. 3ACDF). Notably, and in agreement with [6]’s implementation, the correspondence with the original SCA analysis of the S1A dataset [5] is strong, despite some methodological differences. Namely, [5] performed a SVD, not an ICA (see, Fig. S1A for SVD plots). Residues were also manually assigned to the different sectors (XCoRs here), allowing for the possibility that a residue could belong to multiple sectors. Finally, for each dataset, we observe a strong robustness of the results with respect to the values of the similarity threshold and the gap filtering threshold (see e.g. Fig. S2 for the S1A dataset).

Differences for the DHFR dataset (Fig. 3BE, Fig. S3) are mostly observed in the purple IC and its corresponding XCoR, which includes many residues from the C-terminal region. This region is highly gapped in the original MSA (over 30%, Fig. S3E) and, hence, likely correspond to residues filtered in [25], but not in our analysis. Note also that this XCoR is qualitatively different from the other XCoRs, as it consists of contiguous residues in the primary sequence, in contrast to the scattered residues characteristic of sectors [5].

### XCoR visualization along phylogenetic trees and associated metadata

One of the most salient features of COCOA-Tree is its ability to visualize protein sequences along phylogenetic trees with mapped metadata. When XCoRs are treated as sequence subsets, this greatly facilitates interpretation of the origins of coevolutionary signals. In particular, it supports both qualitative and quantitative assessments of how coevolutionary patterns relate to whole-sequence phylogeny. When enzymatic function is used as metadata, it also allows direct evaluation of how coevolutionary patterns link to protein function – a central aspect of the sector hypothesis of functional decomposition in proteins [5, 6].

To illustrate these capabilities, we first show the results obtained for the red XCoR of the S1A serine protease family, corresponding closely to the red sector of [5] and predicted to determine substrate specificity. Specifically, we display in the same figure: a phylogenetic tree obtained from the full sequences; metadata associated with these sequences, including functional information (catalytic specificity) and taxonomic information; the corresponding XCoR sequence; and a heatmap indicating the level of identity between any two XCoR sequences. A visualization based on 442 sequences is provided in Fig. S4, with a reduced subset shown in Fig. 4A for clarity. From these representations, and more particularly when considering 442 sequences, we first observe that red XCoR sequences do not globally follow the phylogeny of the full sequences: some sequences that are evolutionarily distant cluster tightly at the XCoR level. This corroborates the functional nature of sectors [5, 6]. Yet, local block patterns suggest that phylogeny still contributes to the coevolutionary signal. Second, the red XCoR sequences of chymotrypsins exhibit a divergence pattern distinct from that of the other enzymes. Specifically, they diverge more rapidly than those of the other enzyme families. For example, compared to chymotrypsins, red XCoR sequences of trypsins remain highly similar even across large phylogenetic distances (Fig. 4A, Fig. S4). This suggests that chymotrypsin functionality can be achieved through a broad range of XCoR sequence variants, whereas trypsin functionality appears to be associated with a much more constrained XCoR sequence space.

**Figure 4.**
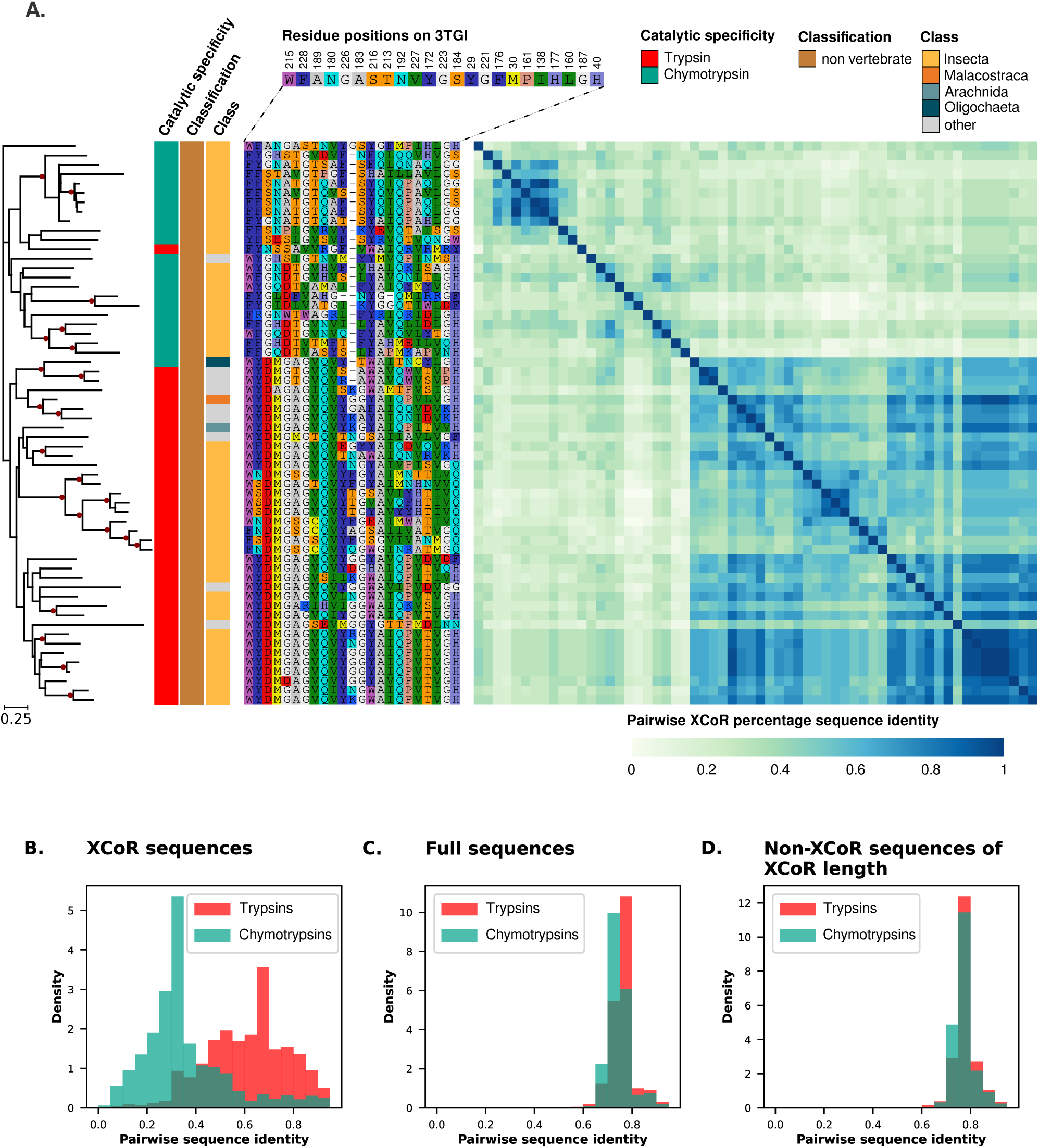
Phylogenetic tree and metadata visualizations of S1A red XCoR (sector). **A.** We constructed a phylogenetic tree using 442 full sequences from the S1A dataset. For clarity, the tree shown here is a subset of 60 sequences (see Methods). From left to right, the following information is provided for each sequence: enzymatic function (catalytic specificity, first column), taxonomic information (next two columns), XCoR sequences sorted by decreasing IC score (with the residue position on the rat trypsin PDB structure [3TGI] indicated at the top), and a heatmap showing the percentage sequence identity between pairs of XCoR sequences. Red dots in the tree represent branches that harbor a high support (UFBoot *≥* 95 %); the scale represents substitutions per site. **B/C/D.** Distribution of identity levels considering trypsin and chymotrypsin enzymes separately between (**B**) pairs of XCoR sequences, (**C**) pairs of full-length sequences and (**D**) pairs of non-XCoR sequences composed of randomly picked residues to match the red XCoR length.

One might question whether these observations simply reflect low confidence in certain nodes of the phylogenetic trees – due to the high divergence of the sequences. We thus used an alternative approach to test their validity. Namely, we calculated the distribution of identity levels across all pairs of XCoR sequences (Fig. 4B) and compared it with the distribution obtained from all pairs of full-length sequences (Fig. 4C) as well as that obtained from random non-XCoR subsequences with the same XCoR length (Fig. 4D), analyzing chymotrypsins and trypsins separately. Our analysis reveals a clear separation between the XCoR identity distributions, with chymotrypsins exhibiting substantially lower identity values (median *≃* 0.3) than for trypsins (median *≃* 0.6). In contrast, the identity distributions of both the full sequences and the non-XCoR sequences largely overlap, with respective median values of *≃* 0.75 and *≃* 0.74 for chymotrypsins, and *≃* 0.74 and *≃* 0.75 for trypsins. Consistent with these results, sequence logos reveal greater diversity for chymotrypsins than for trypsins (Fig. S5).

As a second interesting illustration, we discuss results concerning the blue XCoR, which corresponds closely to the blue sector of [5] that has been experimentally shown to be related to the thermodynamic stability of the enzymes. Using the same tree visualization as for the red XCoR, our results indeed reveal an interesting scenario. First, as already discussed in the original analysis of [5], the coevolutionary signal underlying this XCoR reflects “organism type,” and more specifically, the vertebrate-invertebrate classification. Here, as shown for the full set of 442 sequences (Fig. S6) and, for clarity, for a subset of 83 sequences (Fig. 5), our analysis further reveals that the coevolutionary signal results from a strong conservation of the XCoR sequence in vertebrates, as indicated both by the sequence information of the XCoR and the identity heatmaps. In contrast, the XCoR sequence exhibits poor conservation in invertebrates. In the line of previous critics of the coevolutionary origins of sectors [15], one might thus argue that this sector reflects more of a conservation property (here specific to a subset of organisms) than a coevolutionary trend. It is nevertheless remarkable that this part of the sequence – and specifically this part, as revealed by comparing the distribution of identities in the XCoR (Fig. 5B), in the full sequences (Fig. 5C) and in random non-XCoR subsequences with the same XCoR length (Fig. 5D) by separating vertebrates from invertebrates – appears to be under strong constraints in vertebrates. It is also interesting that coevolutionary analyses allow the detection of this type of behavior.

**Figure 5.**
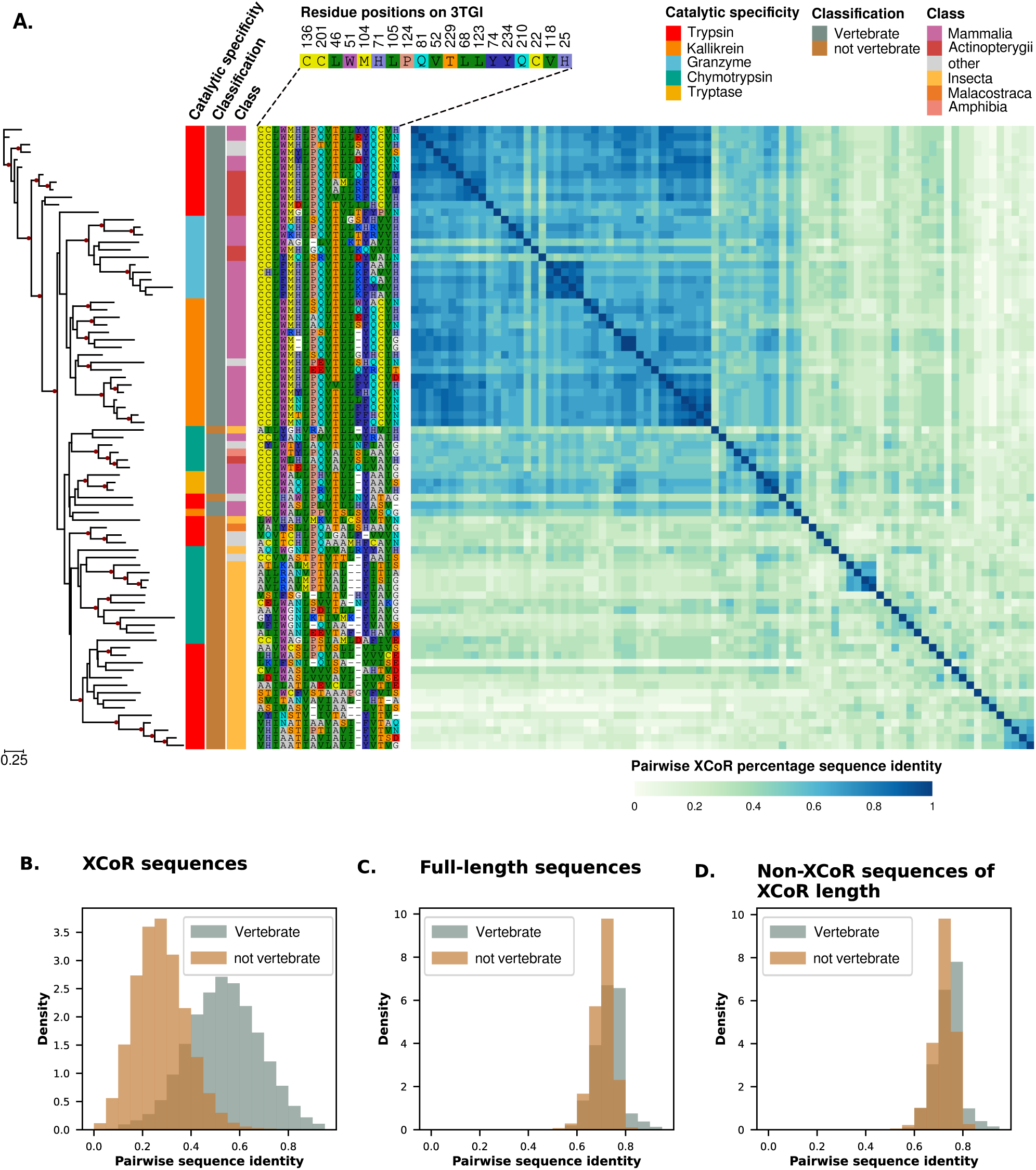
Phylogenetic tree and metadata visualizations of S1A blue XCoR (sector). **A.** As in Fig. 4A, we considered a phylogenetic tree built from the 442 full sequences from the S1A dataset (supp. Fig. S6). For clarity, the tree shown here is built from a subset of 83 sequences, which span a part of the phylogenetic tree that is different from Fig. 4A (see Methods). The reported information is identical to that in Fig. 4A. **B/C/D.** Distribution of identity levels considering vertebrates and invertebrates separately between (**B**) pairs of XCoR sequences, (**C**) pairs of full-length sequences and (**D**) pairs of non-XCoR sequences composed of randomly picked residues to match the blue XCoR length.

Finally, for completeness of the analysis, we provide in Fig. S7 the same visualization for the green XCoR, which corresponds closely to the catalytic-activity-associated green sector. As already discussed in [5], one can observe that this XCoR largely reflects strong conservation (see also the sequence logos in Fig. S8).

### Export of results to PyMOL format

The notion of sectors and, more broadly, of coevolutionary patterns is closely linked to the spatial arrangement of residues within protein folds. A key aspect of interpreting sectors – which have been shown to be composed of residues that form physically connected networks [5, 6, 24] – therefore relies on visualizing these residue groups on the three-dimensional structure of the protein [6]. To facilitate such visualization, COCOA-Tree provides example Python scripts to be used with PyMOL, allowing users to map XCoRs onto any available 3D structure, with residues color-coded according to their contribution to the corresponding IC. As an example, in Fig. S1C we show for the S1A dataset the three XCoRs identified by COCOA-Tree mapped onto the rat trypsin structure.

### Systematic comparison of coevolution metrics

The unified API of COCOA-Tree makes it straightforward to compare the outcomes of coevolution analyses that differ only in the metrics and corrections used to compute coevolution scores. As an example, in Fig. 6 and Fig. S9 (and in Fig. S10 and S11 for the DHFR and Rhomboid datasets), we compare XCoRs obtained for the S1A dataset using three coevolution metrics (SCA, MI, NMI), as well as MI combined with the commonly used APC correction. A central difficulty in fully automating such comparisons is that ICs (and thus XCoRs) have no natural ordering. To compare results, we therefore performed a global clustering of all XCoRs under the constraint that XCoRs obtained with the same method are not clustered together. This allows us to evaluate how well XCoRs match across methods. To that end, each XCoR from each method is ultimately assigned one of three colors: red, green, or blue. Two remarks are then necessary to clarify the comparison. First, for consistency, we considered exactly three XCoRs per method, although the number of relevant ICs is in principle method-dependent (see Decomposition in the Approach section). Second, for methods other than SCA, the colors do not imply any a priori correspondence with the original SCA sector colors. For instance, the red XCoRs from MI-related methods match each other but share no residues with the red SCA XCoR, indicating that they correspond to a completely different unit. In contrast, and interestingly, the green XCoRs obtained with NMI and MI+APC (but not MI) closely match the green SCA XCoR, while the blue XCoRs from MI and MI+APC (not NMI) share a subset of residues with the blue SCA XCoR. Finally, it is worth noting that the MI-related red XCoR stands out for its remarkable stability across the various MI-based metrics and correction schemes. A detailed analysis (Methods) reveals that this XCoR comprises 23 contiguous positions located at the N-terminus of the proteins and corresponds to a highly unstructured signal peptide (supp. Fig. S12), consistent with previous findings on the extracellular serine protease of *Staphylococcus epidermidis* [29]. Similar to the purple XCoR observed in the DHFR dataset, this XCoR is thus qualitatively distinct from sector-like XCoRs, which are instead typically scattered along the primary sequence of the proteins.

**Figure 6.**
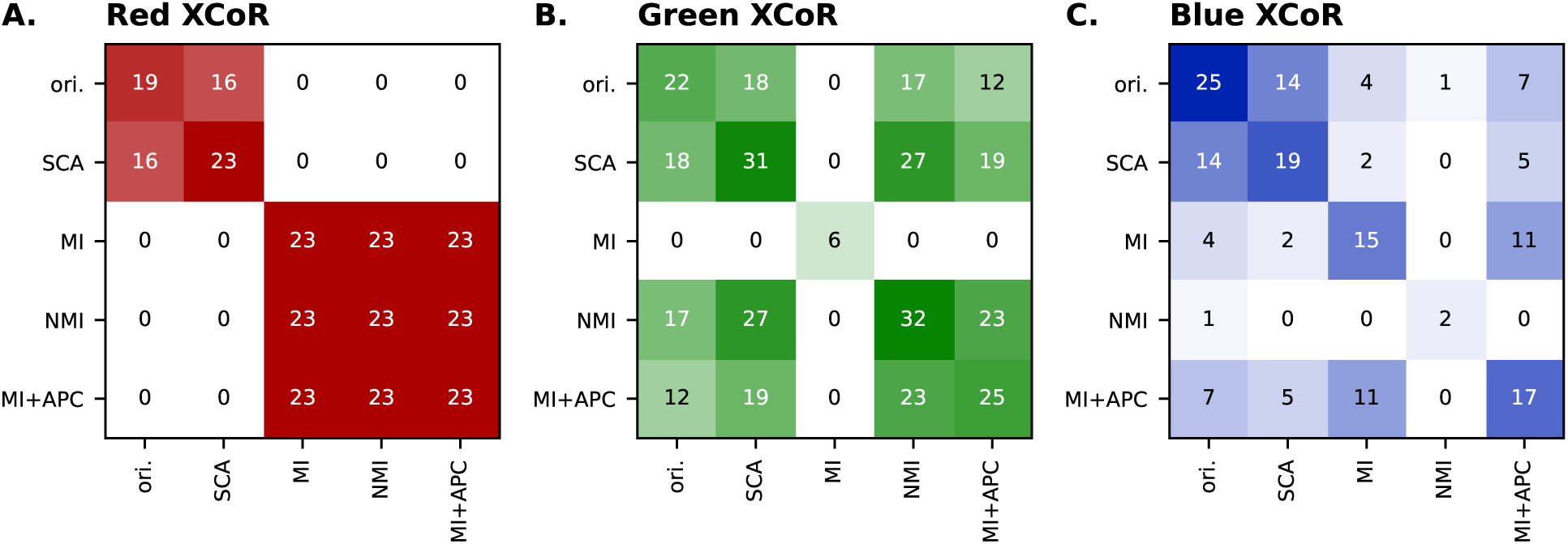
Comparing different coevolution metrics with COCOA-Tree on the S1A dataset. Overlap in residues for **A.** the red XCoR, **B.** the green XCoR, **C.** the blue XCoR between the original results of [5] (Halabi), COCOA-Tree using the SCA matrix (SCA), mutual information (MI), normalized mutual information (NMI), and mutual information corrected with APC (MI+APC). The diagonal of each matrix corresponds to the number of residues found for each method in each XCoR. Off-diagonal numbers correspond to common residues found between XCoRs of two different methods. The full confusion matrix is provided in Fig. S9.

As confirmed with the DHFR and Rhomboid datasets (Fig. S10 and S11), and consistent with previous discussion [14], these results show that different coevolution metrics can yield qualitatively different coevolutionary units. This should not come as a surprise: MI-based metrics tend to emphasize variable positions, since MI increases with information content, whereas SCA emphasizes conserved positions through its weighting scheme [5]. This weighting has been criticized in cases where a single sector emerges, suggesting that sectors in this case may simply correspond to conserved positions [15]. Yet, we find that NMI and MI+APC sometimes find XCoRs that substantially overlap with SCA-derived ones. To further understand these observations, we report in Fig. 7 for the S1A dataset, residue conservation (Shannon entropy) versus the cumulative coevolution score, with residues colored by XCoR. Overall, one can observe that SCA identifies highly conserved positions, MI highlights variable ones, and NMI and MI+APC capture residues spanning both extremes. This explains why NMI and MI+APC sometimes identify XCoRs that are very similar to those detected by SCA. Consistently, similar patterns to the S1A dataset are observed for the DHFR and Rhomboid datasets (Fig. S13 and S14), with a substential overlap of at least two XCoRs obtained by SCA on the one hand, and two XCoRs obtained by MI and MI+APC on the other.

**Figure 7.**
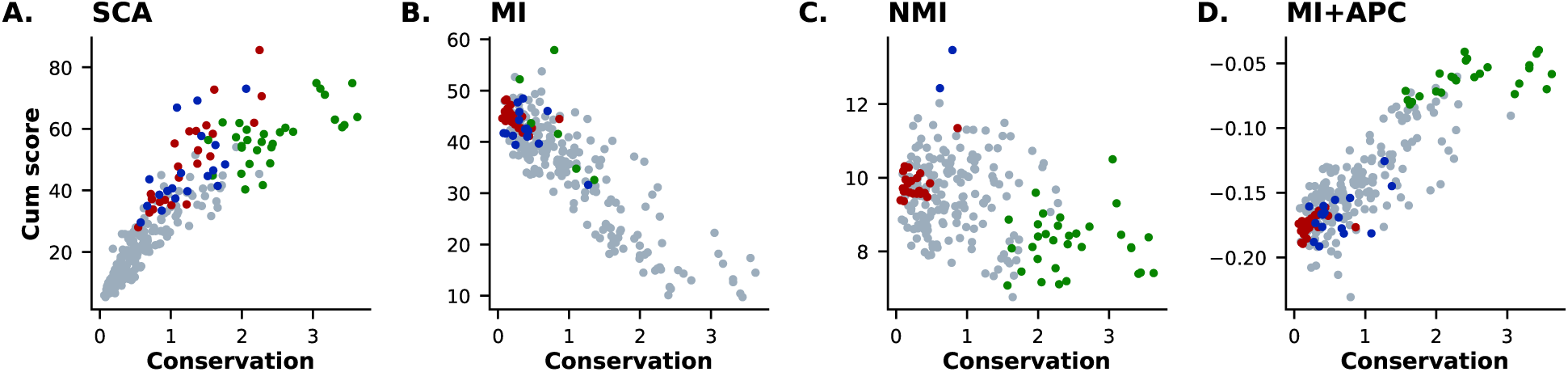
Conservation level of XCoRs identified by different metrics in the S1A dataset. In each panel, corresponding to a given coevolution metric and correction, each point represents a residue, plotted as its cumulative score (the sum of the corresponding row or column in the coevolution matrix) versus its conservation level (measured as the Shannon entropy of its amino-acid composition). Colors indicate residues assigned to XCoRs by COCOA-Tree for each coevolution metric.

### Highlighting coevolutionary patterns in DCA analysis

Although COCOA-Tree does not include a module for performing DCA analyses, the library can be used to visualize the results of such analyses in relation to the phylogeny of input sequences and metadata. As an example, we present some results in the context of the S1A dataset, which we subjected to an adabmDCA analysis [30], an adaptive Boltzmann-machine learning framework for DCA [22] — see Methods for details. Specifically, we selected ten top DCA pairs and used the COCOA-Tree library to plot them on the phylogenetic tree of the S1A dataset, along with functional information (catalytic specificity) and taxonomic information (Fig. S15). In general, for most pairs, we observe that certain coordinated variations in amino acid composition coincide with phylogenetic changes, but these effects do not necessarily occur at the same location or depth in the tree (Fig. S15).

Fig. 8 further highlights some interesting coevolutionary patterns by showing the results for three specific pairs (136-201, 189-226, and 57-195 on the rat trypsin PDB structure [3TGI]) at two locations in the phylogenetic tree of the sequences — indicated by dashed rectangles in Fig. S15. The signal of pair 136-201 is strongly associated with the vertebrate versus invertebrate classification, with a disulfide bridge (CC amino acid pair) that is highly conserved in vertebrates (Fig. 8A) and varies significantly in invertebrates (Fig. 8B). Interestingly, the disulfide bridge is also occasionally observed in invertebrates. Pair 189-226 presents an interesting case of a variation that follows neither phylogeny nor taxonomic classification but rather the functional classification of enzymes, particularly between trypsin and chymotrypsin. Moreover, we can observe a recurrent pattern of compositional swap (DG composition exchanged for GD). These observations are in accord with the known implication of residue 189 in the conversion between trypsin and chymotrypsin [31]. Also, this pair, which belongs to the red sector, has been identified as a “secton”, an intermediate unit between (DCA-like) structural contacts and large functional (sector-like) units [32]. Finally, pair 57-195 involves a highly conserved residue pair for which the coevolutionary signal results from a coordinated variation in composition for only a few sequences that are distant in the tree.

**Figure 8.**
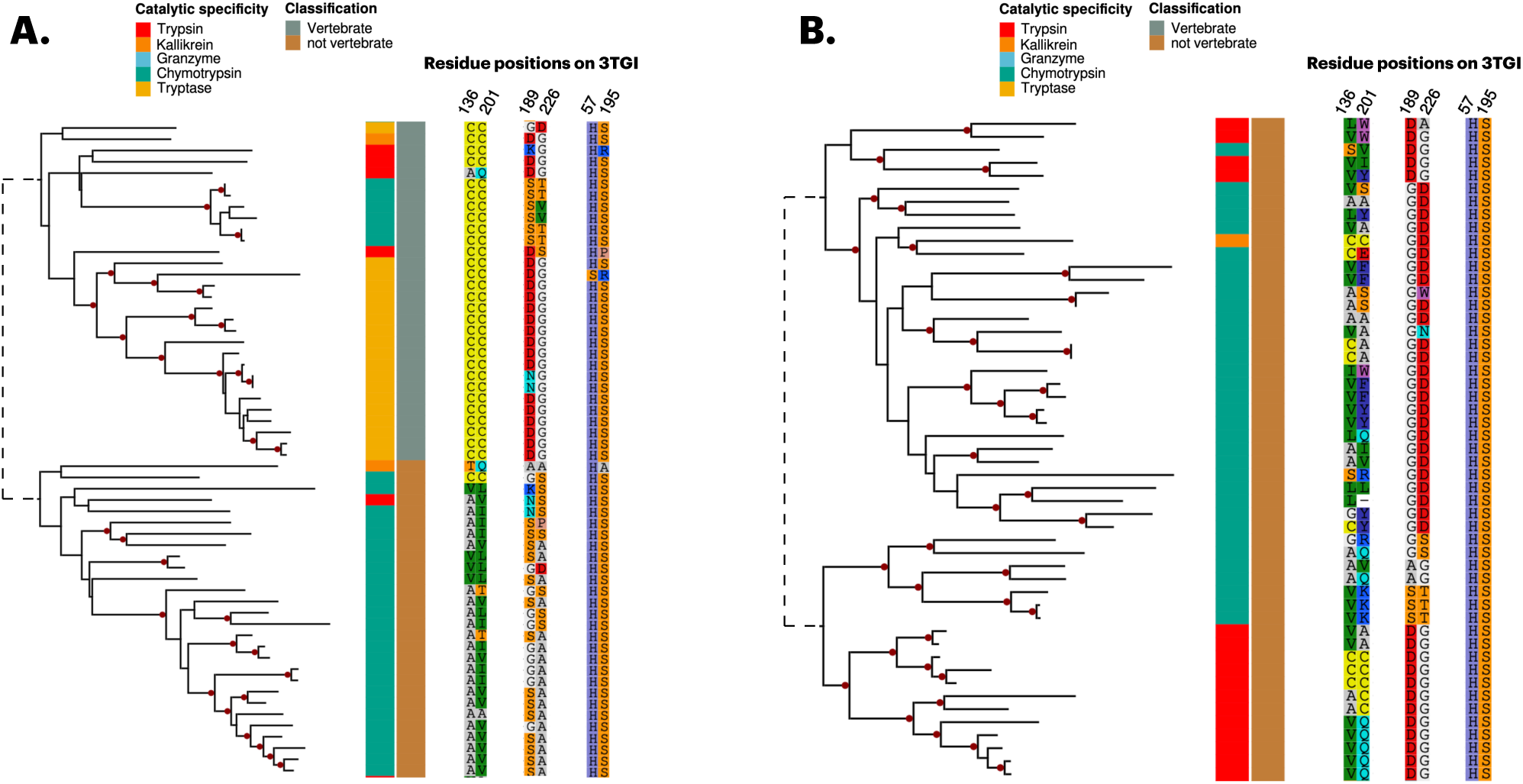
Phylogenetic tree visualization of DCA coupling pairs extracted from the S1A dataset. With the goal of highlighting various co-evolutionary patterns, we selected three DCA pairs (positions 136-201, 189-226 and 57-195 on the rat trypsin PDB structure [3TGI]) and two tree subsets (A and B) from the entire tree (Fig. S15). Regarding metadata, for the sake of clarity, we only show functional and classification annotations.

In summary, in addition to a relatively heterogeneous impact of phylogeny, several types of coevolutionary patterns can be distinguished for DCA pairs that are independent of phylogeny, in accord with the poor influence of phylogeny on the strongest DCA couplings [16, 21]. Visualization in the context of functional metadata also allows to readily discriminate between patterns associated with changes in function from those expected to be associated with structural constraints.

## Conclusion

We have presented COCOA-Tree, a Python library designed to systematize co-evolutionary analyses and visualize their results with phylogenetic trees. By confronting the outcomes of dimensionality reduction methods with those derived from the reconstruction of sequence evolutionary history, our aim is twofold: (1) to better understand the origin of correlations identified by the former methods and (2) to gain deeper insights into the evolutionary trajectories of sequences. To demonstrate these capabilities, we reanalyzed three enzyme families previously studied and discussed in the context of SCA. The interest in considering SCA-related works is twofold. First, the most recent implementation of SCA [6] relies on a dimensionality reduction method (ICA) that is highly generic and agnostic to the co-evolution metric used. This flexibility underpins COCOA-Tree’s ability to equivalently generate and compare results from different metrics. Second, SCA predictions have been poorly discussed in the context of the phylogeny of the underlying sequences, making it an approach that is sometimes difficult to reconcile with phylogenetic methods [1]. By integrating SCA results with phylogenetic approaches, we thus not only demonstrate the potential of COCOA-Tree but also deepen our understanding of the co-evolutionary signals that give rise to the sectors discussed in the literature. Note, finally, that by restricting our analysis to specific published datasets, we did not explore the potential impact of certain variables that could influence the outcome and interpretation of the results. Such variables include the size and diversity of the input sequences, the quality of the input MSA, the number of retained ICs, as well as the various filters applied to exclude residues or construct the phylogenetic trees. Addressing these aspects is beyond the scope of this article, although performing such analyses would be facilitated by the library.

The case of the serine protease family, as discussed here as examples, provides a clear illustration of the COCOA-Tree potential. Namely, we identified two previously unnoticed yet crucial properties that we believe will enhance our understanding of the evolution of these enzymes and, more specifically, of their sectors. First, we revealed a qualitative difference in the conservation of the XCoR (sector) predicted to govern substrate specificity, for enzymes that differ precisely in their substrate. Specifically, the red XCoR (sector) is significantly less conserved in chymotrypsins than in trypsins, particularly at large phylogenetic distances. From the perspective of this sector, the functional sequence space for chymotrypsins thus appears broader than that for trypsins. It is then worth mentioning that synthetic conversions have only been reported from trypsin to chymotrypsin [31], while the reverse conversion has not [33]. We premise this reflects different selective constraints acting on the composition of the sector. Second, we revealed a qualitative difference in the conservation of the XCoR predicted to govern the thermodynamic stability of enzymes, depending on the type of organism to which the enzyme belongs. Specifically, the blue XCoR (sector) is highly conserved in vertebrates but variable in invertebrates. This observation leads to two implications. The first is a prediction: [5] showed that mutations in the blue sector of rat trypsins (a vertebrate) qualitatively altered the enzyme’s melting temperature without affecting its catalytic activity. We predict much milder – or possibly negligible – effects in the case of invertebrate trypsins. The second implication concerns the nature of the co-evolutionary signal associated with this sector. It is indeed legitimate to question whether it is appropriate to speak of coevolution in this case, as the signal arises from the conservation of the sector within a specific clade rather than from simultaneous or coordinated sequence changes. This ties into broader discussions about the nature of sectors [15] and the molecular origins of correlations in residue substitutions [16, 34], which may sometimes (or often) be indistinguishable from phylogenetic relationships. While this question remains actively studied (see, e.g., [19, 35]), we believe that our tool, COCOA-Tree, will be particularly valuable in advancing our understanding of these issues.

An additional interesting feature of COCOA-Tree is the ability to compute standard coevolution metrics and corrections natively. As shown above, this makes it straightforward to compare results across methods and is therefore expected to improve the prediction of potential determinants of enzyme functionalities. We also remind that COCOA-Tree allows users to implement their own coevolution metrics in a straightforward manner.

Finally, COCOA-Tree can be used to display groups of residues of interest independently of the residue-detection methods included in the library. This option becomes particularly interesting in the context of DCA. Indeed, to our knowledge, there is no simple-to-use library available to visualize a list of residue pairs in the context of phylogenetic trees and associated metadata. By providing access to the code that allowed us to generate Fig. 8 and Fig. S15 (see script availability), users can easily produce figures for their own data. Here, we have shown that such use of COCOA-Tree allows for a relatively straightforward distinction of signals associated with phylogeny, functional and contact constraints in DCA pairs. Along the same line, COCOA-Tree could be used to explore more deeply the impact of methods that have been proposed to mitigate phylogenetic contributions [13, 21]. For instance, one could visualize and compare, on the phylogenetic tree, pairs or groups of residues identified with and without these methods. Altogether, COCOA-Tree should thus also help distinguish structural, functional, and phylogeny-induced couplings between residues.

## Methods and datasets

### PySCA

Part of the COCOA-Tree code is derived from the PySCA repository available at https://github.com/ranganathanlab/pySCA. This includes more specifically the estimation of the number of relevant ICs, the attribution of a residue to a specific XCoR, and the function to compute ICA. The corresponding methods are detailed in [6].

### Datasets provided as part of COCOA-Tree

#### S1A serine proteases

The S1A dataset consists of enzymes that catalyze peptide bond hydrolysis through a conserved mechanism, with substrates (*i.e.*, targeted proteins) that vary across its members. Three sectors (named red, green, and blue), corresponding to the first three ICA components, have been identified and, based on both structural and experimental evidence, have been linked to specific functional roles within the protease family [5]. Namely, the red sector surrounds the S1 pocket, which was shown to be associated with the substrate specificity of these enzymes [31]; that is, this sector is expected to “control” which proteins are degraded by a specific enzyme. Next, the green sector comprises residues directly involved in the chemical reaction catalyzed by the enzyme and is therefore believed to control the catalytic core of the protease family. Finally, mutations in the blue sector have been shown to be associated with the thermodynamic stability of the proteins and is believed to control the local stability of protein regions that may be associated with other functions.

#### Dihydrofolate reductases (DHFRs)

DHFR catalyzes the reduction of dihydrofolate to tetrahydrofolate, a key step in folate metabolism and nucleotide synthesis. Beyond its therapeutic relevance as a target to prevent cell proliferation, DHFR also serves as a model system for latent allostery [24]. Namely, using SCA, [24] identified a sector, composed of several XCoRs, that links the active site to distal surface regions acting as hot spots for the emergence of allosteric control. Recently, [25] combined SCA with molecular dynamics simulations and showed that coevolving residues also exhibit correlated motions. Notably, in this study, four XCoRs were analyzed as distinct units rather than merged into a single sector as in [24], two being located around the enzyme active site.

#### Rhomboid proteases

Just like S1A serine proteases, the widely distributed membrane superfamily of rhomboid proteases degrades proteins by catalyzing the hydrolysis of peptide bonds. However, in contrast with the S1A superfamily, the serine protease active site of rhomboid proteases is membrane-embedded and formed from different transmembrane segments. A SCA analysis of this family revealed three significant ICA components, two of which were fused due to their high level of correlation, leading to the proposal of two sectors, named Sector1 and Sector2 [26]. Analogous to the red and blue sectors in the S1A family, Sector1 and Sector2 are thought to determine, respectively, the substrate specificity of the family and the structural stability of the protein fold and functional sites. In particular, the relevance of Sector1 for controlling substrate specificity was experimentally demonstrated by converting the substrate specificity of one enzyme into that of another simply by grafting the sequence of its Sector1 [26].

### Coevolution metrics and corrections

The following section presents the equations used to compute the coevolution metrics and corrections used in COCOA-Tree. We consider an alignment *S* of *M* sequences and *L* positions with 20 amino acids + gaps. The binary array *x_si_^a^* equals 1 if sequence *s* has amino acid *a* at position *i* and 0 otherwise.

#### Statistical Coupling Analysis coevolution matrix

Here, we recall the mathematical definition of the SCA coevolution metrics as presented in [6]. First, a covariance-like matrix is computed considering all pairs of positions *i* and *j* and all pairs of amino acids *a* and *b* at these positions:

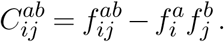

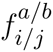 stands for the frequency of amino acid *a/b* at position *i/j* and 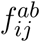 for the joint frequency of amino acids *a* and *b* at positions *i* and *j*. These frequencies are computed as:

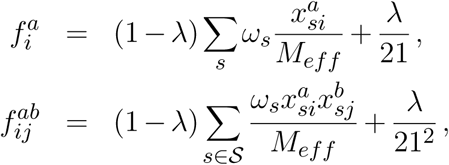

where: 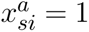 if amino acid *a* is found at position *i* in the sequence *s*, 0 otherwise; *ω_s_* are the sequence weights; *M_eff_* = ∑*_s_ω_s_* is the number of effective sequences; and *λ* a regularization (pseudo-count) to account for the finite size of the MSA. This covariance-like matrix is then weighted by a conservation measure of amino acids at each position, i.e., 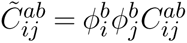 with:

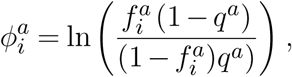

where *q^a^* is a background distribution of amino acid [6]. Finally, the SCA co-evolution matrix is computed as the Frobenius norm of 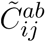:

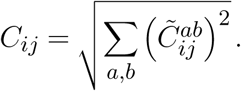

#### Mutual information

The matrix of mutual information (MI) is computed as:

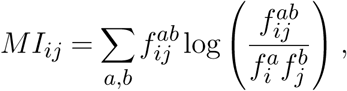

with 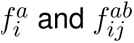 computed as in SCA.

#### Normalized mutual information

The normalized mutual information (NMI) consists in normalizing the mutual information by the joint Shannon entropy 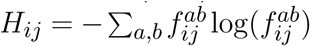. The NMI coevolution metrics thus reads:

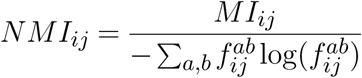

#### Average product correction

The average product correction (APC) between positions *i* and *j* as defined by [8] is computed as

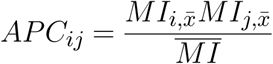

where 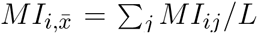 is the mean mutual information involving position *i* and 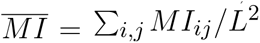 is the overall average mutual information over all pairs of positions. The matrix of mutual information corrected by APC is then given by:

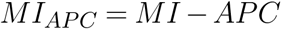

#### Entropy correction

Although not investigated here, to simplify the presentation of the library, COCOA-Tree includes an entropy correction as defined by [12], computed as:

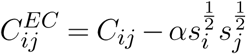

where the strength of the correction, *α*, is given by 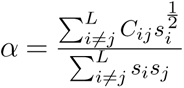.

### Phylogenetic analyses

The tree visualizations of Fig. S4, Fig. S6, and Fig. S7 were generated considering 442 sequences of the original MSA (S1A dataset) for which both functional and taxonomic information were available. The resulting MSA was trimmed with ClipKIT [36] using the kpic-smart-gap option. Maximum-likelihood phylogenetic trees were computed using the IQ-Tree software, version 2.0.3 [37]. UFBoot supports were computed with 1000 replicates and the best model was assessed for each alignment using the default parameters. In Fig. 4 and respectively Fig. 5, we sampled two regions of this tree: (1) a region containing trypsin and chymotrypsin sequences from non vertebrate organisms (delimitated with a black square on Fig. S4), within which 60 sequences were randomly sampled, and (2) 83 sequences from vertebrate and non vertebrate organisms. To obtain robust trees, because the original MSA was trimmed relative to rat’s trypsin sequence [5], we re-extracted the full sequences from the NCBI database (https://www.ncbi.nlm.nih.gov/). Both subsets of sequences were then aligned using MAFFT-DASH [38] (structural alignement) and the resulting alignments were trimmed with BMGE [39] using the default parameters and a BLOSUM30 matrix. Phylogenetic trees were then computed using the IQ-Tree software with the same parameters as mentioned earlier.

### Protein logos

To plot protein logos, we computed the frequency of each amino acid for each residue for all sequences belonging to the group of interest. We then plotted logos with LogoMaker [40].

### Structural and functional analysis of the MI-based red XCoR (S1A dataset)

The MI-based red XCoR comprises 23 contiguous positions located at the N-terminus of proteins, from position 107 to 129 in the unfiltered MSA. To further investigate this set of positions, we inferred an AlphaFold structure for a sequence in which the sector contains no gaps (51871601, trypsin 10 precursor from *Rattus norvegicus*). The N-terminal region corresponds to a low-confidence portion of the AlphaFold model and appears highly unstructured, suggesting the presence of a signal peptide (Fig. S12). Consistently, analysis with SignalP 6.0 [41] shows that 73% of proteins containing at least one residue from this XCoR are annotated as having a signal peptide.

### Extraction of DCA pairs in the S1A dataset

We inferred the DCA pairs using the adabmDCA code version 2.0 [30]. To this end, we tested different L2 regularization parameters, with *λ ∈ {*0.0001, 0.001, 0.01, 0.1*}*. Following the recommended procedure, a DCA score was computed by imposing a zero-sum gauge on the couplings, using the Frobenius norm, and applying ACP (Averaged Product Correction). We also discarded pairs of residues whose distance along the linear sequence was fewer than 5 amino acids. In this context, the top 10 pairs with the highest DCA scores were found to be the same for all four regularization parameters, although not in the same order.

## Acknowledgements

We thank Hugo Mutschler and Morgane Roger-Margueritat for valuable feedback on COCOA-Tree, as well as Olivier Rivoire for insightful discussions and feedback on the manuscript.

## Fundings

We acknowledge funding by the French National Research Agency (Project Deepen, ANR-19-CE45-0013-02), by the Initiative of Research in Grenoble-Alpes (IRGA Project GenUbi), and by the French National Research Agency in the framework of the Investissements d’Avenir program (ANR-15-IDEX-02) through the funding of the “Origin of Life” project of the Grenoble-Alpes University.

## Data, script, code, and supplementary information availability

The COCOA-Tree website can be found at https://tree-timc.github.io/cocoatree, and the code at https://github.com/TrEE-TIMC/cocoatree. The code used to reproduce the results and generate the figures can be found at https://github.com/TrEE-TIMC/cocoatree_ms_figure.

## Supplementary Information

## Supplementary Note 1

### Supplementary tables

**Table S1.** Relationship between sectors or ICs and XCoRs. We map each of XCoRs to their original sectors by the best match

| COCOA-Tree | Serine protease | Rhomboid | DHFR |
| --- | --- | --- | --- |
| Red | Red | Cyan | Blue |
| Green | Green | Yellow | Yellow |
| Blue | Blue | Red | Green |
| Purple | . | . | Red |

## Supplementary Note 2

### Supplementary figures

**Figure S1.**
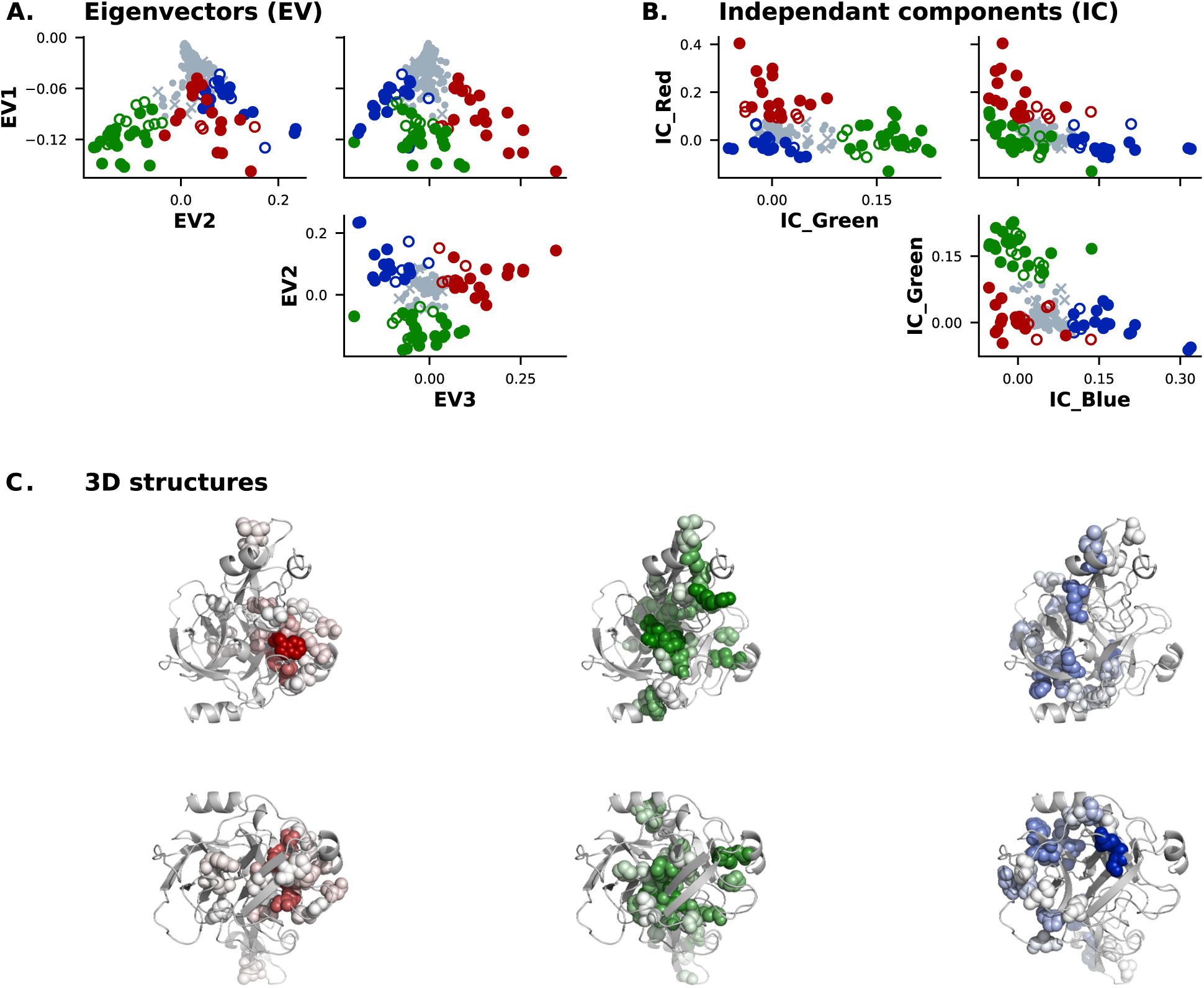
Statistical Coupling Analysis on S1A dataset. A/B. Eigenvectors (A) and ICA decomposition (B) of the SCA coevolution matrix showing the three XCoRs identified by [5] (sectors), those identified by our study (full colored dots), and the XCoR residues only identified by our study (empty colored dots). C. XCoR projections on rat’s trypsin 3D PDB structure [3TGI]. Color intensity increases with the residue’s contribution to the corresponding IC.

**Figure S2.**
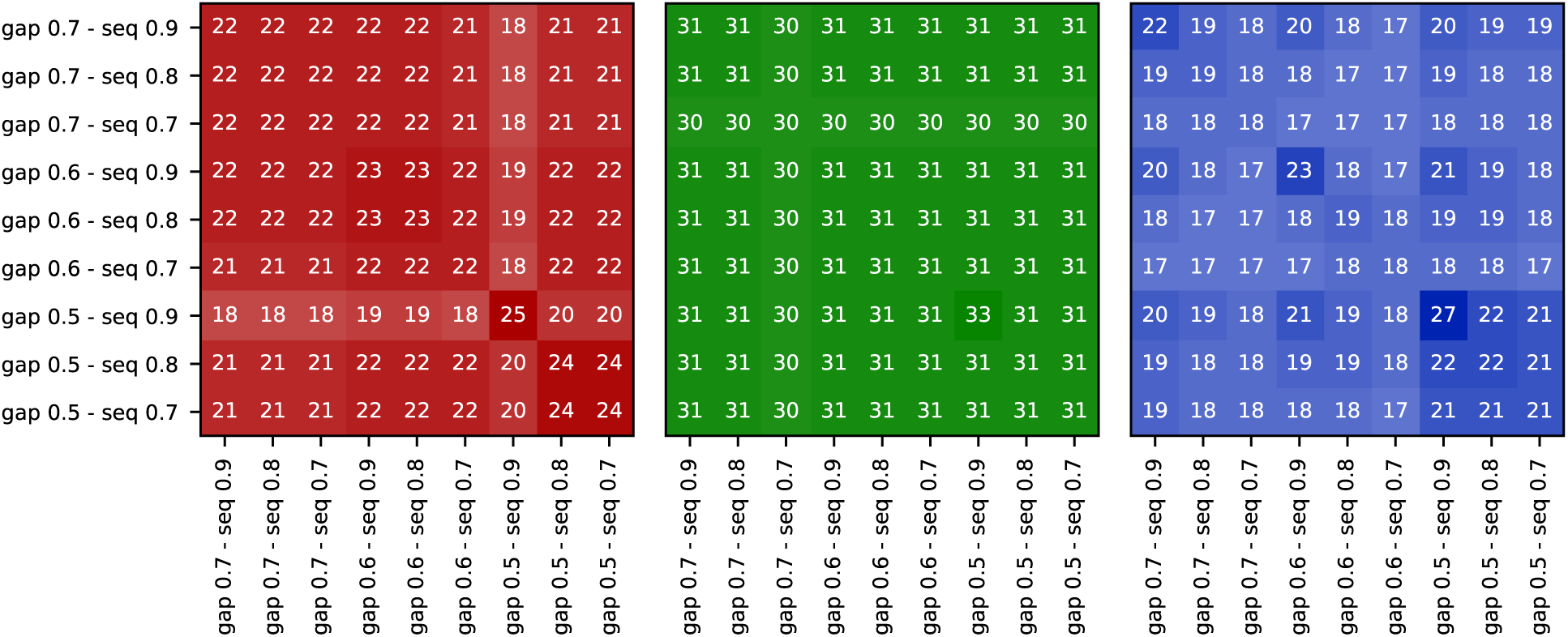
Stability analysis of XCoRs on the S1A dataset varying gap and sequence similarity thresholds. The heat maps show the number of overlaps in residues for the red, green and blue XCoRs obtained from the SCA analysis of the S1A dataset, using values of the gap filtering threshold from 0.3 to 0.5, and of the similarity threshold from 0.7 to 0.9.

**Figure S3.**
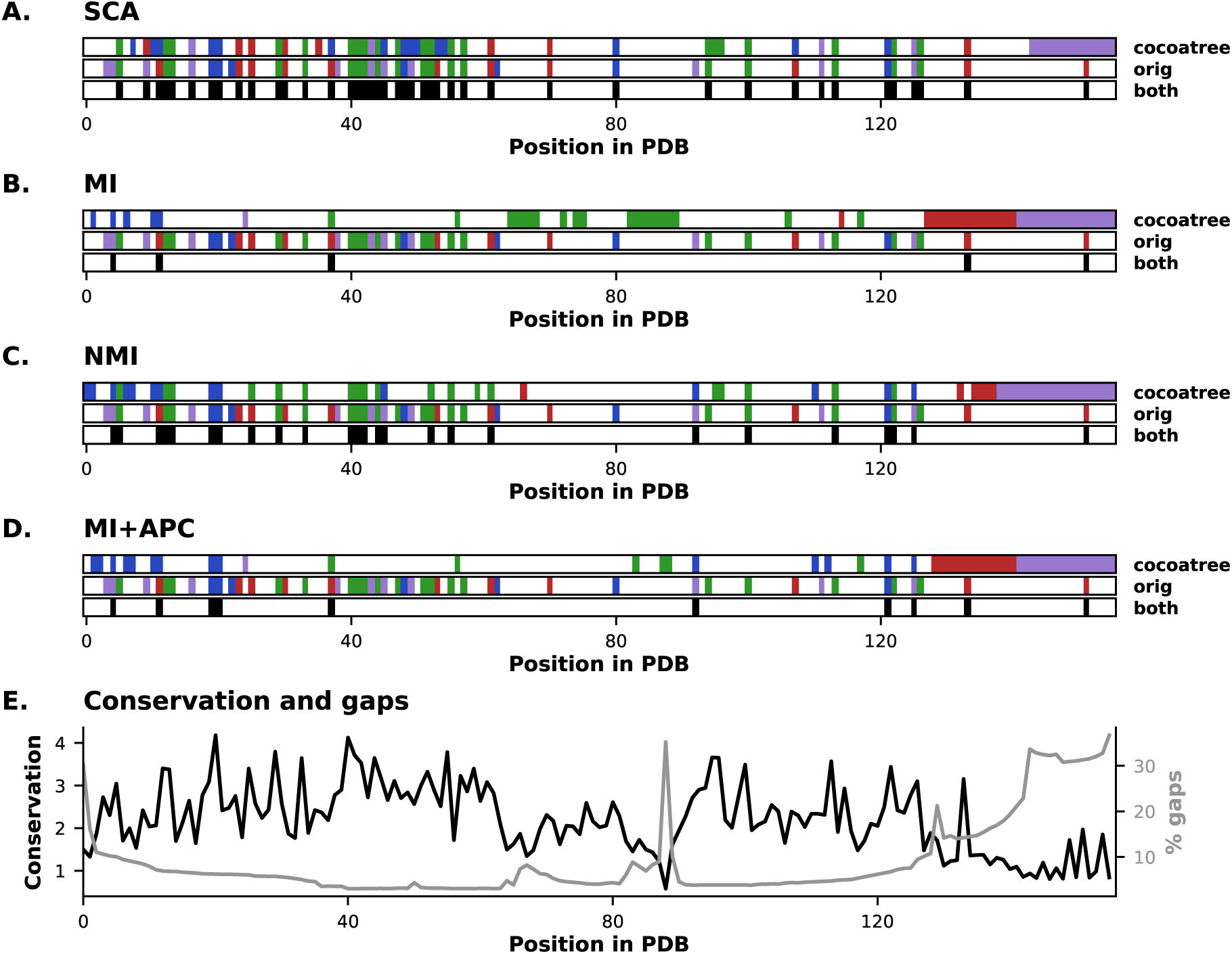
Comparison of COCOA-Tree’s results for the DHFR dataset along *Escherichia coli*’s 3QL3 PDB sequence. A-D. Residue positions that are detected in a XCoR with COCOA-Tree, in the original study, and in both, with SCA metric (A), MI metric (B), NMI metric (C), and MI metric with APC correction (D). E. Conservation and percentage of gaps in the MSA mapped on the PDB sequence.

**Figure S4.**
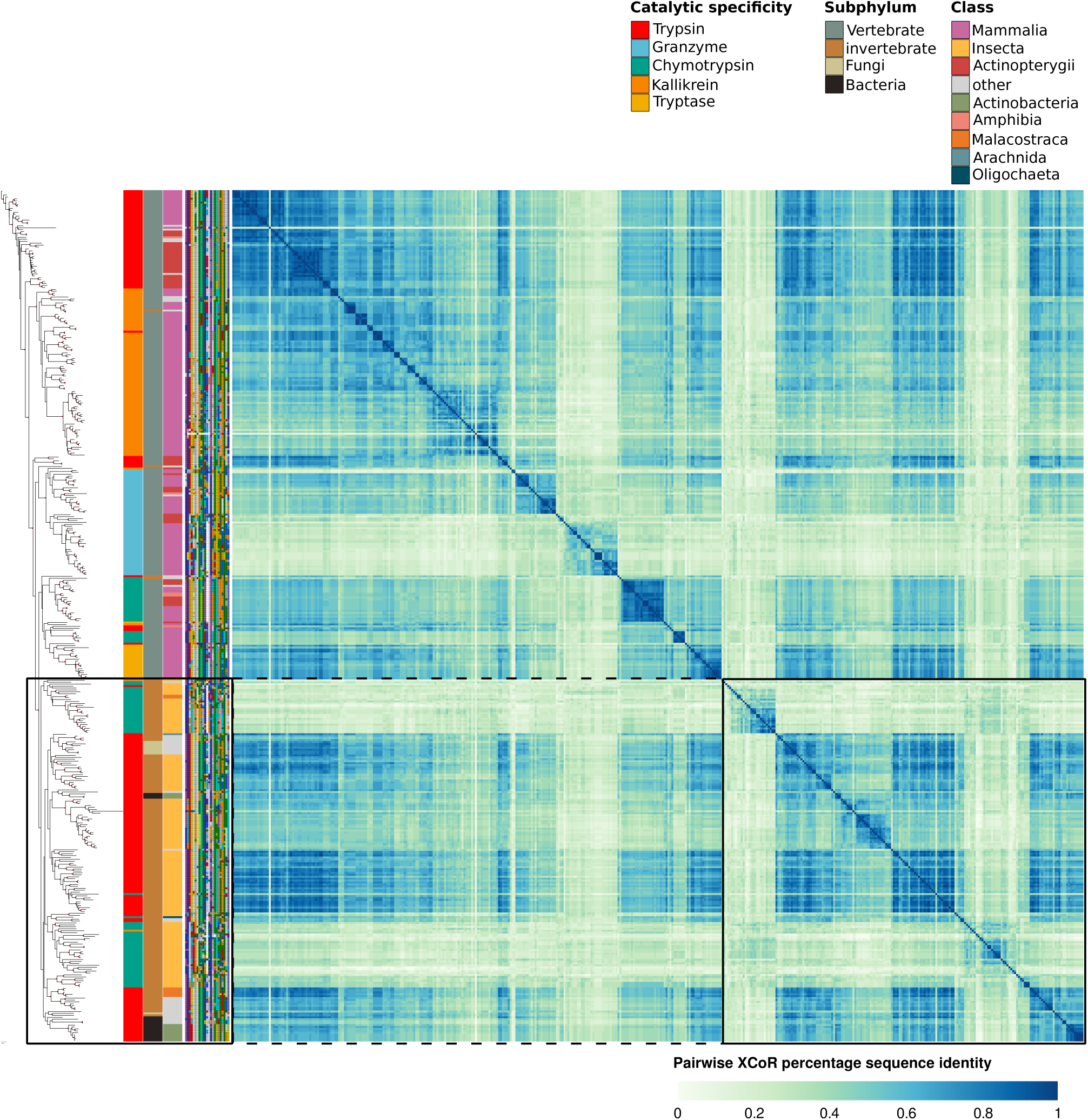
Visualizing the red XCoR of the S1A dataset on a phylogenetic tree (SCA metric, no correction). The tree subset, which is detailed in the main text, is delimitated with black squares. From left to right: a phylogenetic tree computed using 442 sequences from the full S1A serine proteases dataset (red points correspond to 95% and above bootstrap values); sequence annotations (Catalytic specificity, Subphylum and Class) corresponding to each sequence; residue sequence corresponding to the red XCoR (ordered by decreasing contribution to the IC); a matrix of sequence identity computed from the XCoR sequences.

**Figure S5.**
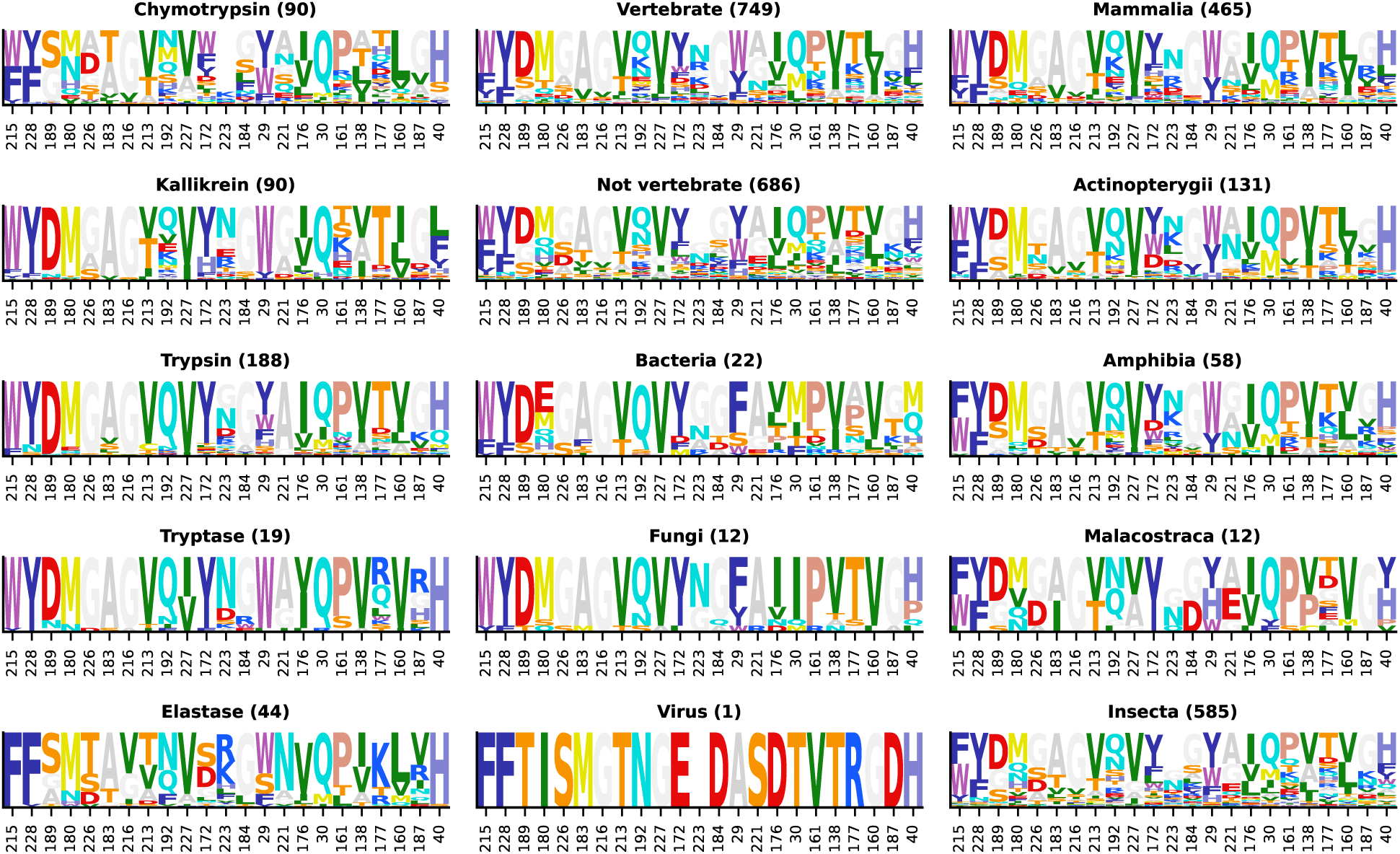
Protein Logo for the red XCoR, grouping sequences by protein specificity (first column), subphylum (second column), and class (third column). Each group’s number of sequences is provided between parentheses.

**Figure S6.**
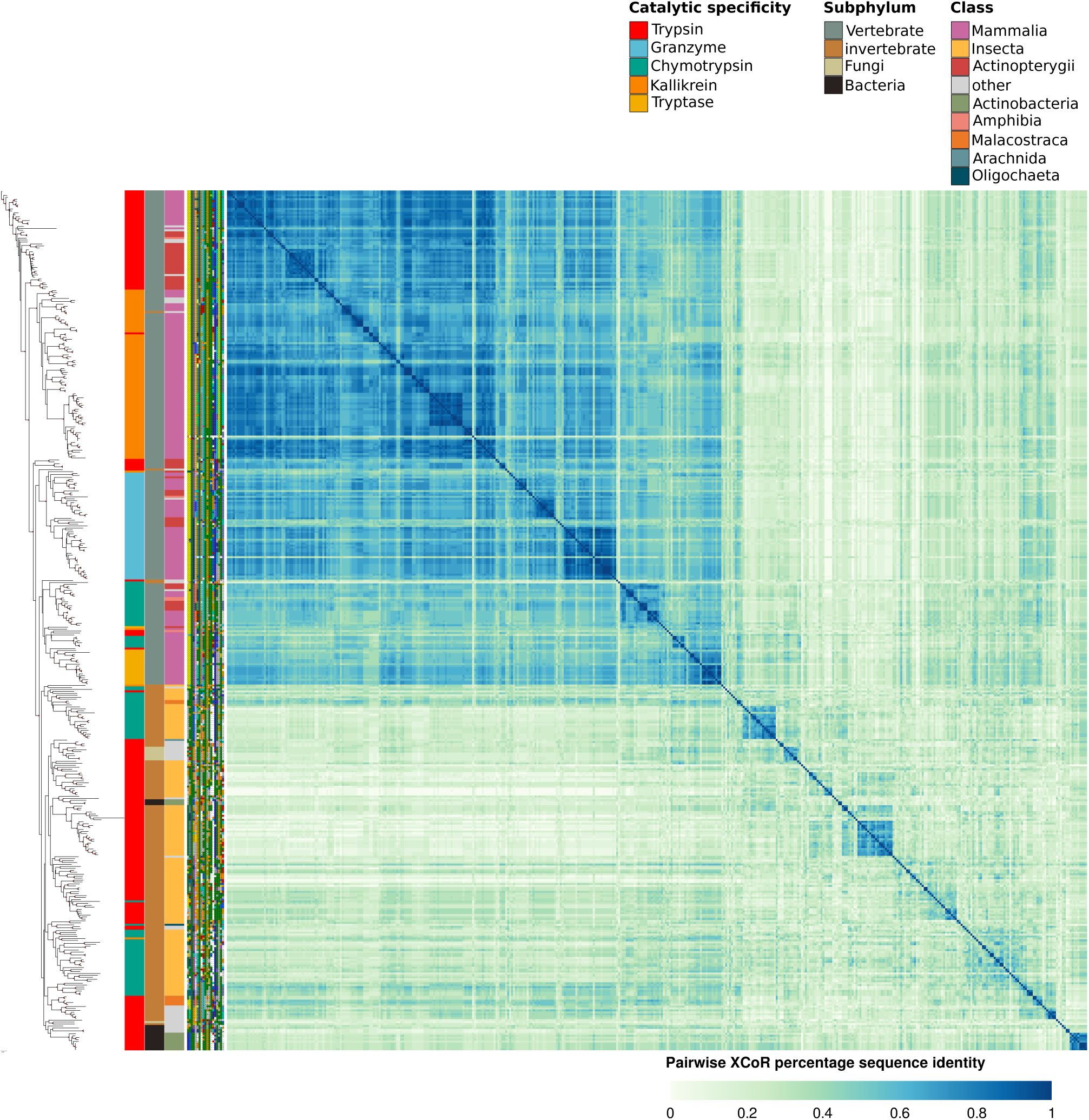
Visualizing the blue XCoR of the S1A dataset on a phylogenetic tree (SCA metric, no correction)

**Figure S7.**
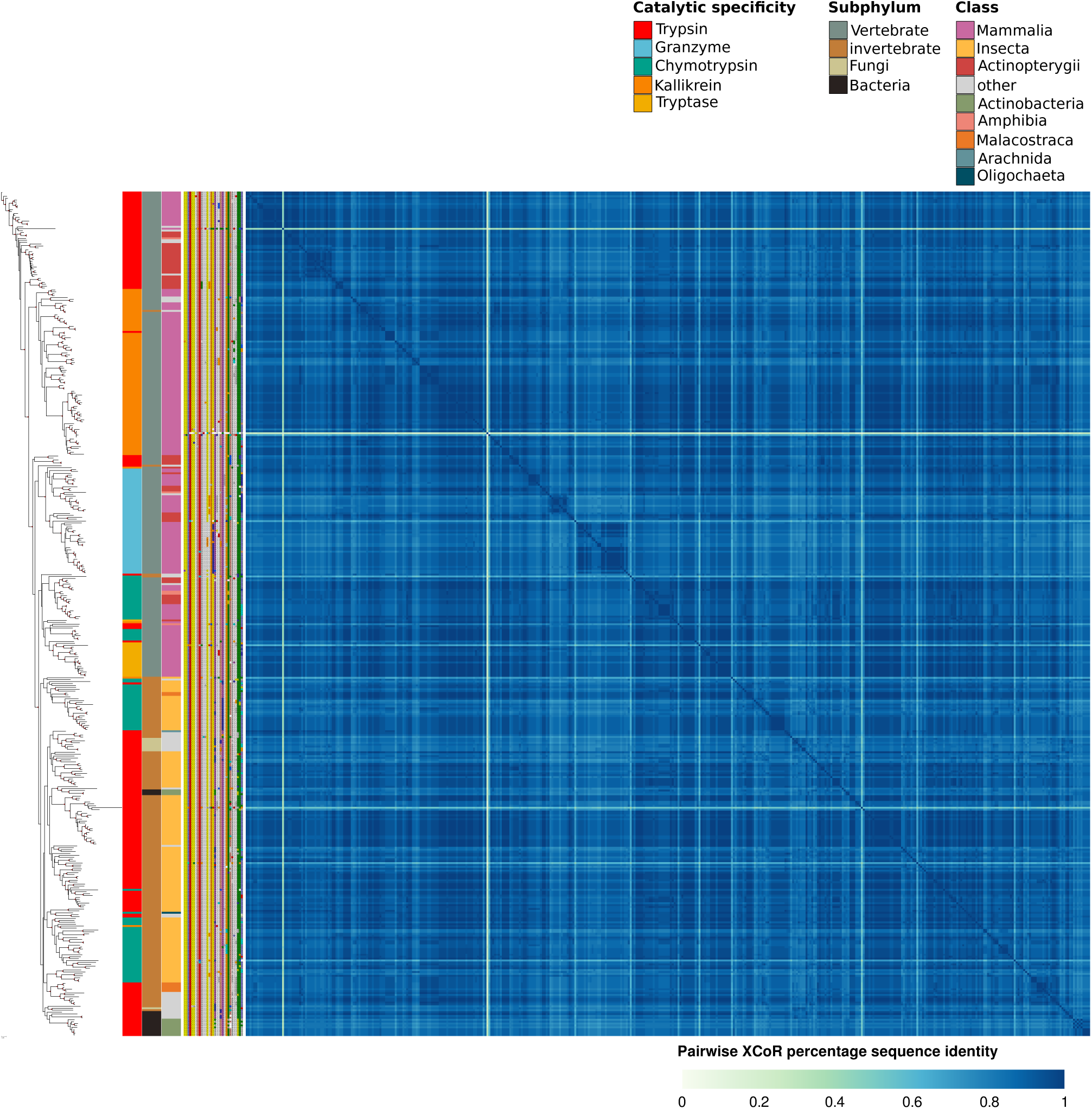
Visualizing the green XCoR of the S1A dataset on a phylogenetic tree (SCA metric, no correction)

**Figure S8.**
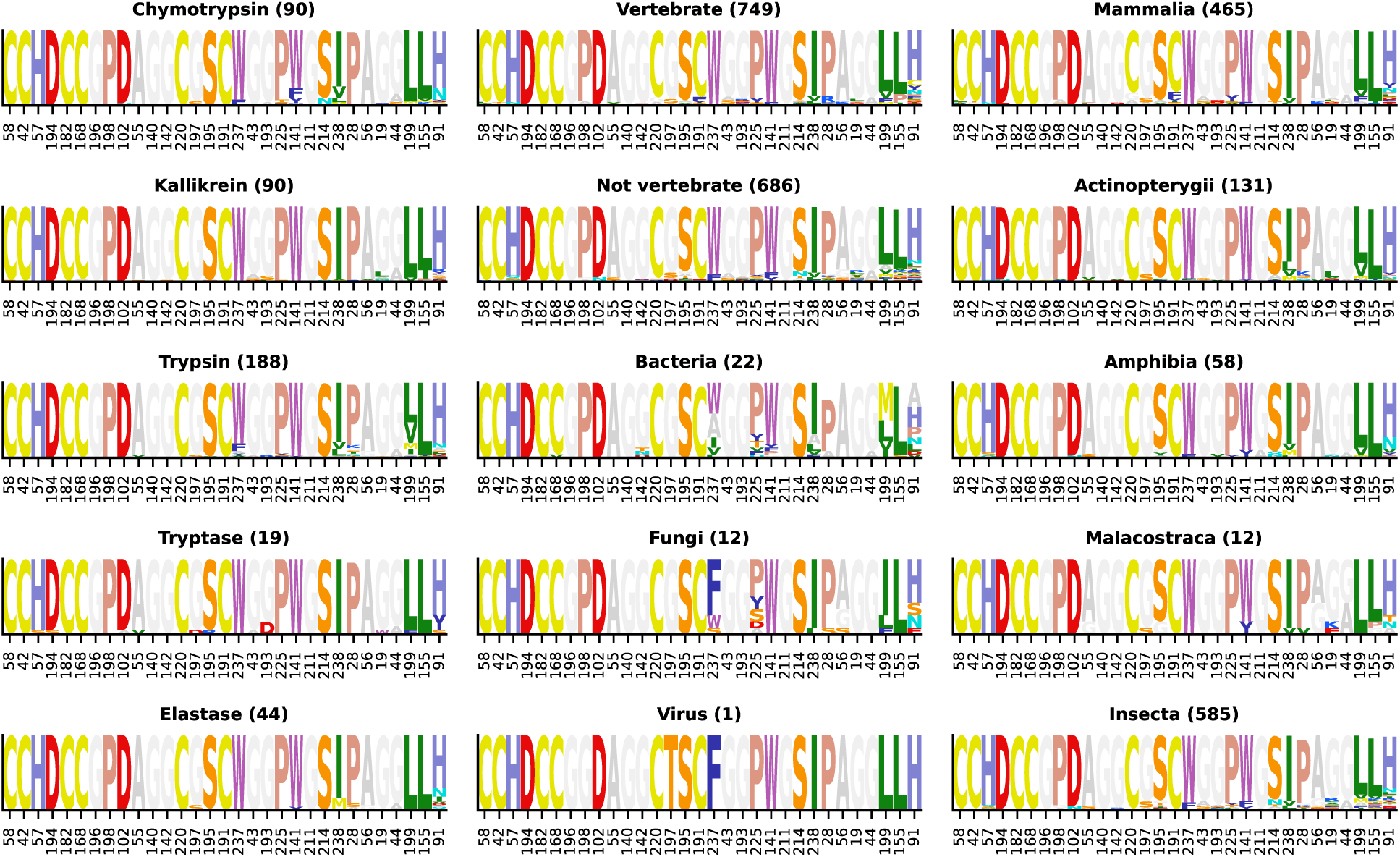
Protein Logo for the green XCoR, grouping sequences by protein specificity (first column), subphylum (second column), and class (third column). Each group’s number of sequences is provided between parentheses.

**Figure S9.**
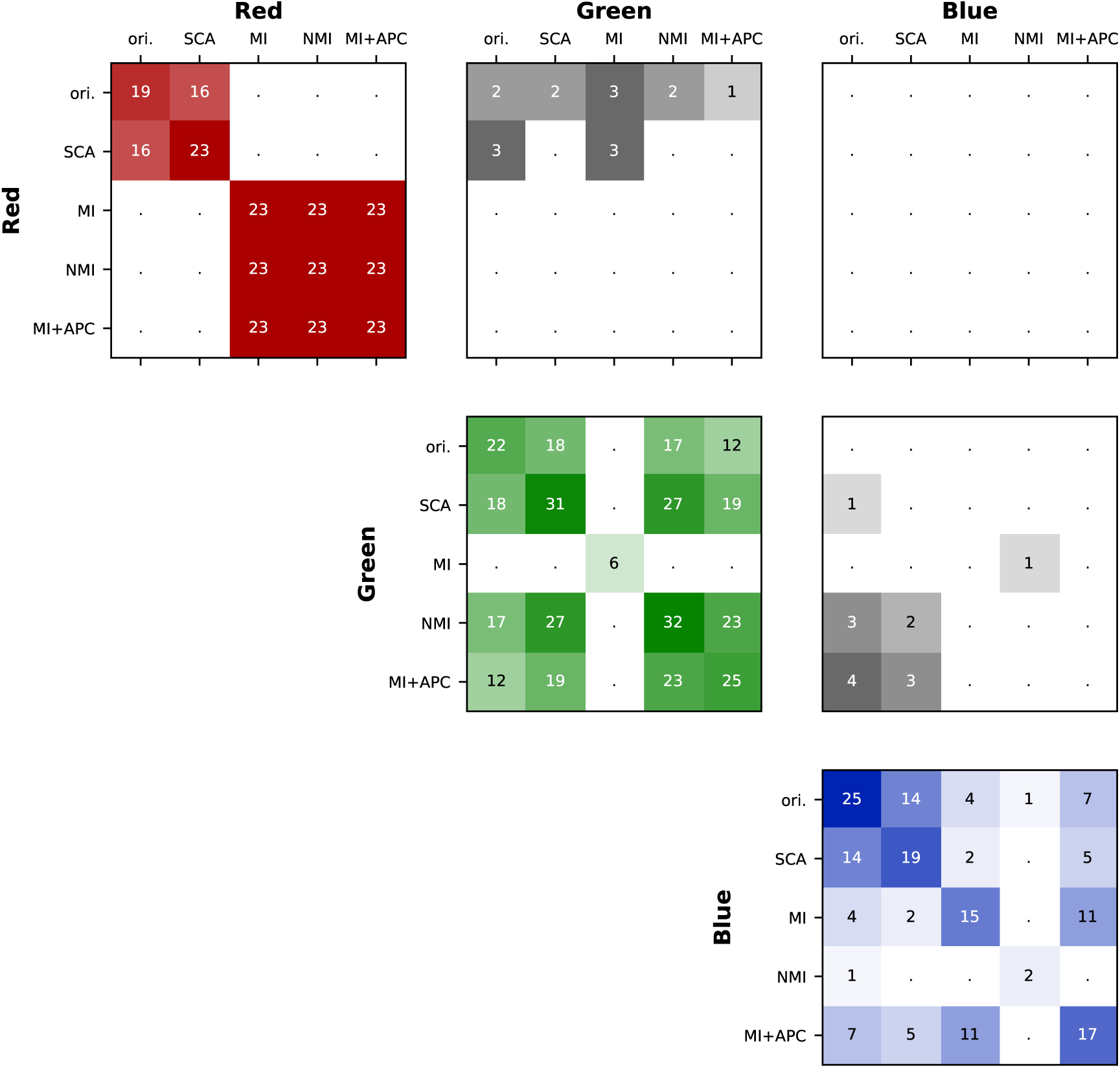
Overlap in sector positions found for the S1A dataset.

**Figure S10.**
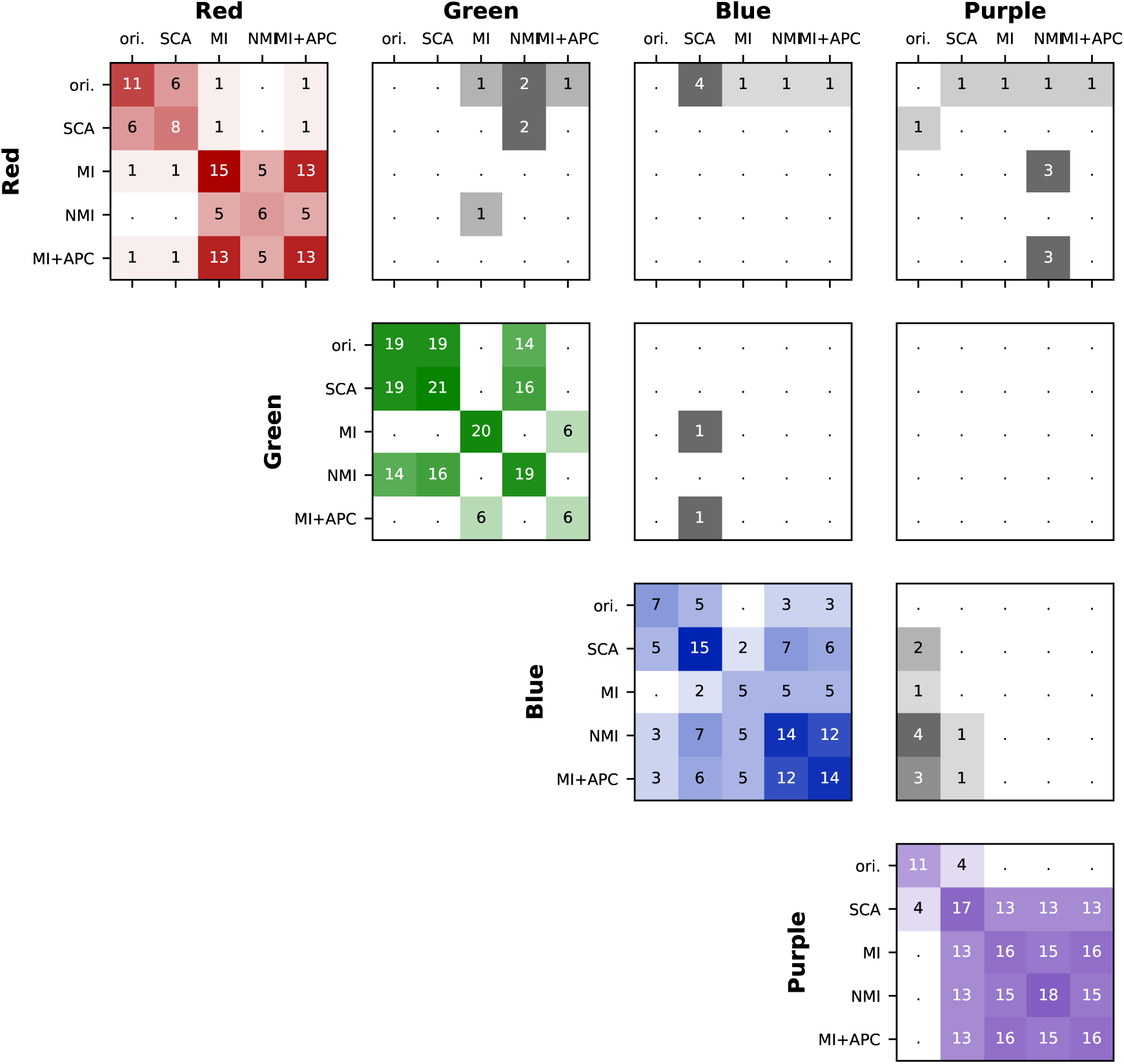
Overlap in sector positions found for the DHFR dataset.

**Figure S11.**
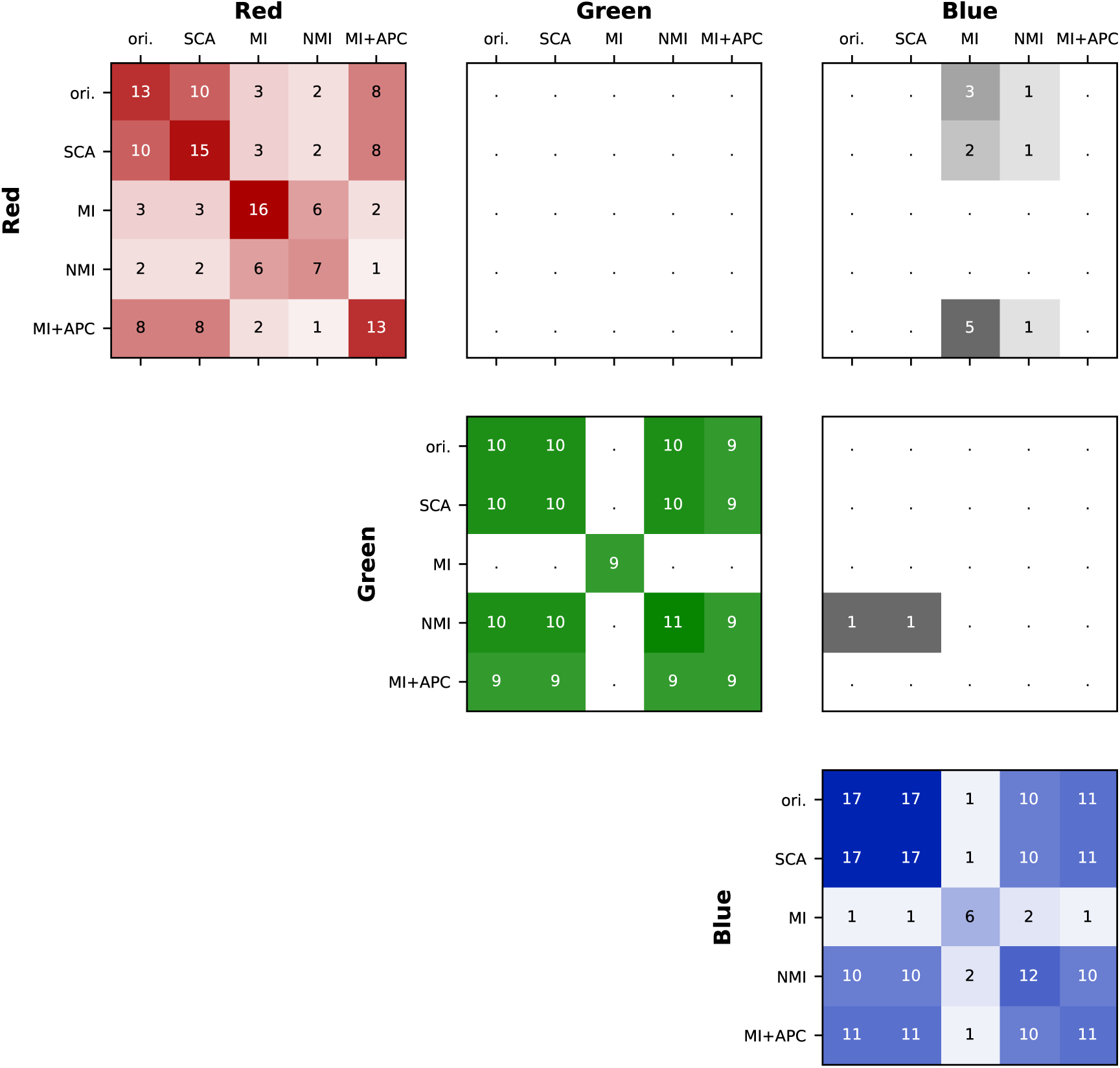
Overlap in XCoR positions found for the Rhomboid dataset.

**Figure S12.**
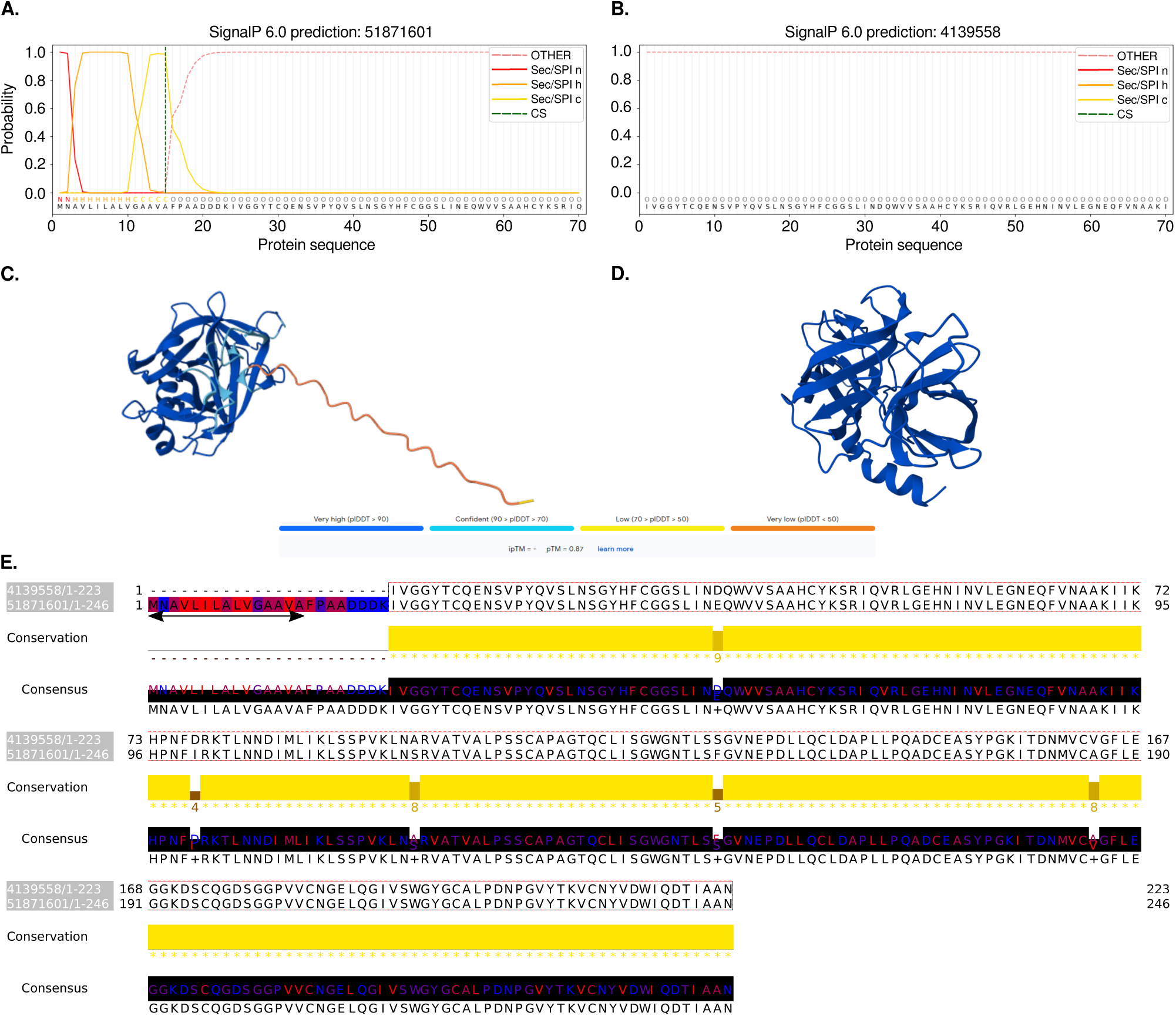
Analysis of signal peptides on two rat’s trypsin sequences. A. SignalP results on rat’s trypsin 10 precursor and B. on rat’s trypsin (3TGI sequence). Each line represents the probability for a residue to belong to a signal peptide (SP) or not (OTHER), and whether it is positively charged (n), hydrophobic (h), or neutral (c); C. AlphaFold prediction based on rat’s trypsin 10 precursor sequence; D. AlphaFold structure prediction based on rat’s trypsin 3TGI sequence. Color represents confidence of the AlphaFold model; E. Jalview visualization of the alignment of rat’s trypsin 10 precursor and 3TGI sequences. The colored residues correspond to the XCoR identified by MI and colors show amino acid hydrophobicity (red hydrophobic, blue hydrophylic). The horizontal double arrow delimitates the signal peptide predicted by SignalP.

**Figure S13.**
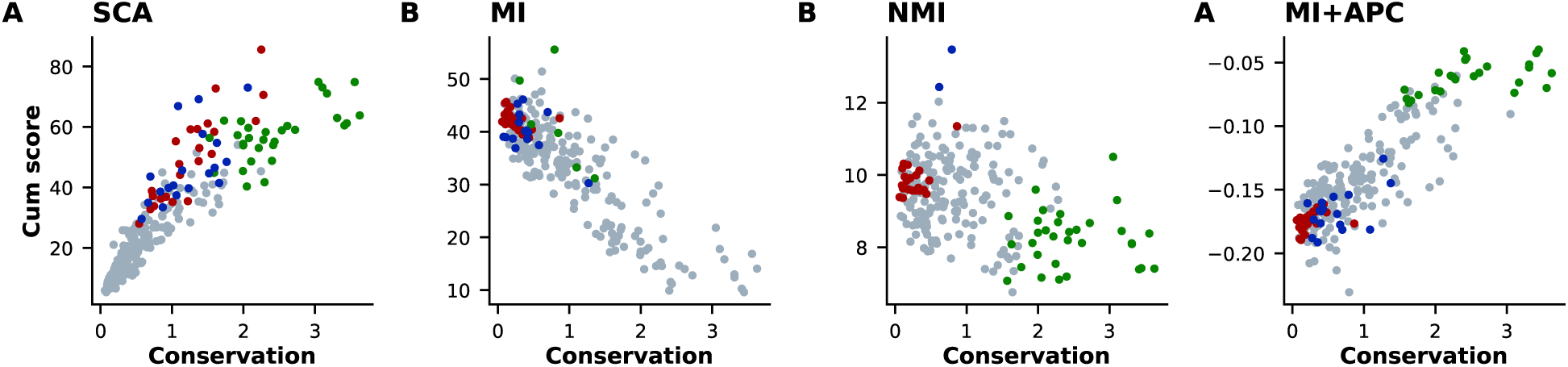
Conservation vs. cumulative score for the DHFR dataset.

**Figure S14.**
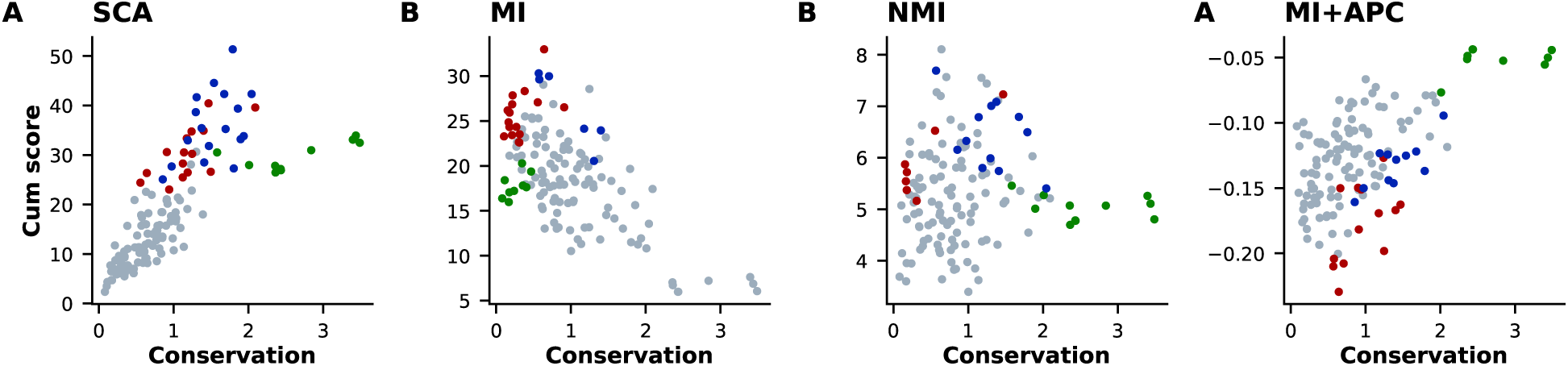
Conservation vs. cumulative score for the Rhomboid dataset.

**Figure S15.**
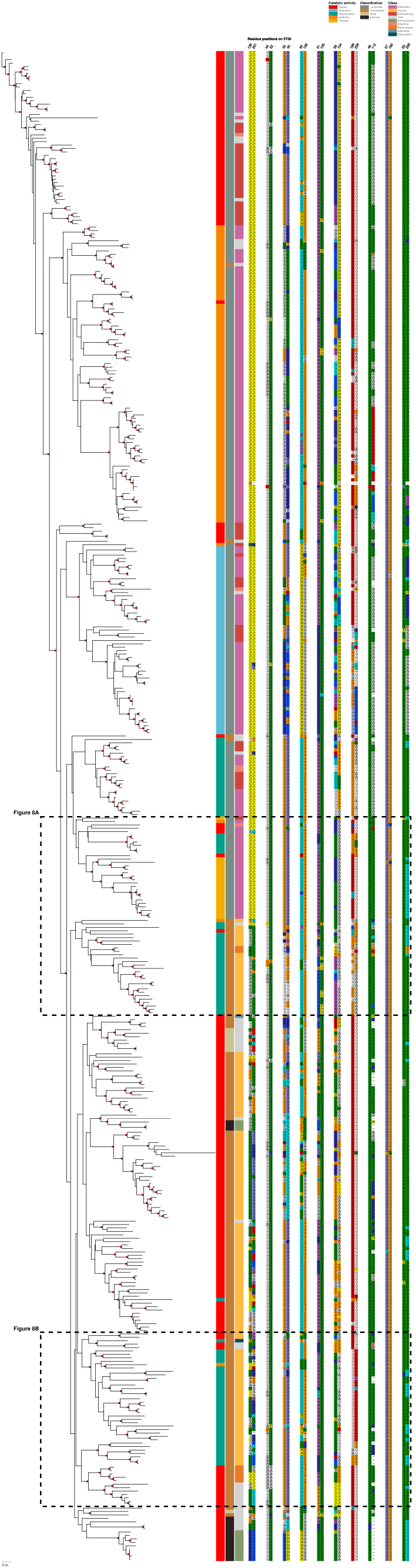
Phylogenetic tree visualization of ten top DCA coupling pairs extracted from the S1A dataset. The tree subset and sequence annotations are identical to the one used to visualize XCoRs – see e.g. Fig. S4. The dashed squares indicate the sequences and pairs that are selected in Fig. 8.

## Footnotes

1 ICA was originally developed in the context of blind source separation, a machine learning task that consists of separating mixed sound signals to recover each independent source [23]. It has since been applied to many types of data, including coevolution metrics, in order to decompose coevolution tendencies into groups that are as statistically independent as possible, that is, leading to possible residual correlations between components [6].

## Notes

https://tree-timc.github.io/cocoatree

https://github.com/TrEE-TIMC/cocoatree

https://github.com/TrEE-TIMC/cocoatree_ms_figure

### Competing Interest Statement

The authors have declared no competing interest.

### Summary of Updates

Our article has been peer-reviewed and recommended by PCI Math Comp Biol. We added the PCI Math Comp Biol badge on the first page. The recommendation can be found at https://doi.org/10.24072/pci.mcb.100684

